# Global profiling of myristoylation in *Toxoplasma gondii* reveals key roles for lipidation in CDPK1 and MIC7 function

**DOI:** 10.1101/719062

**Authors:** Malgorzata Broncel, Caia Dominicus, Alexander Hunt, Bethan Wallbank, Stefania Federico, Joanna Young, Moritz Treeck

## Abstract

*N*-myristoylation is a ubiquitous class of protein lipidation across eukaryotes and *N*-myristoyl transferase has been proposed as an attractive drug target in several pathogens. Functionally the myristate often primes for subsequent palmitoylation and stable membrane attachment, however, growing evidence also suggests additional regulatory roles for myristoylation on proteins. Here we describe the first global chemoproteomic screening of protein myristoylation in *Toxoplasma gondii*. Through quantitative mass spectrometry coupled with validated chemoproteomic tools, we identify 65 myristoylated proteins. We report functionally important myristoylation on the key signalling protein CDPK1 and, surprisingly, myristoylation of the microneme protein 7 (MIC7), a predicted type-I-transmembrane protein. We demonstrate that myristoylation of MIC7 is not important for the trafficking to micronemes, but appears to play a role in host cell invasion. This dataset represents a large fraction of the parasite’s myristoylated proteome and a prerequisite to investigate this modification in *Toxoplasma.*

## Introduction

Toxoplasmosis is affecting approximately one third of the world’s population (Robert-Gangneux and Darde, 2012). It is caused by the obligate protozoan parasite *Toxoplasma gondii* originating from the phylum Apicomplexa. While the majority of human infections are asymptomatic, the disease manifests its severity in immunocompromised individuals, such as those receiving chemotherapy, transplants or HIV/AIDS patients. Key steps in the successful propagation of *T. gondii* infection in the acute phase are orchestrated cycles of invasion and egress from the host cells (Black and Boothroyd, 2000). These crucial processes are regulated by several post-translational modifications (P™s), such as phosphorylation (Gaji et al., 2015; Jacot and Soldati-Favre, 2012; Lourido et al., 2010; Lourido et al., 2012; Treeck et al., 2014), ubiquitination (Silmon de Monerri et al., 2015), and also protein lipidation, such as palmitoylation and myristoylation (Alonso et al., 2012; Frenal et al., 2014).

While the extent of protein palmitoylation in *Toxoplasma* has been investigated (Caballero et al., 2016; Foe et al., 2015), the myristoylated proteome remains uncharacterised. Myristoylation can prime proteins for subsequent palmitoylation and stable protein-membrane association, it has also been shown to facilitate protein-protein interactions (PPIs) and affect protein structure and stability (Martin et al., 2011; Wright et al., 2010). *N*-myristoylation is an irreversible, predominantly co-translational covalent addition of myristic acid to an *N*-terminal glycine (Boutin, 1997; Gordon et al., 1991). It is catalysed by *N*-myristoyl transferase (NMT), which is conserved in *Toxoplasma* and has been reported to be a prominent drug target in fungal (Devadas et al., 1995; Nagarajan et al., 1997), *Trypanosome* (Frearson et al., 2010; Wright et al., 2016) and *Leishmania* infections (Hutton et al., 2014; Wright et al., 2015). In *Plasmodium falciparum* (the causative agent of malaria), inhibition of NMT leads to severe pleiotropic consequences affecting parasite development (Schlott et al., 2019; Wright et al., 2014), highlighting the importance of myristoylation for pathogen survival and progression.

An *N*-terminal ‘MG’ motif is a requirement, but not a predictor of myristoylation. Circa 6% of all gene products in *Toxoplasma* contain the *N*-terminal glycine and an in silico prediction of myristoylation suggests that ∼ 1.8% of all *T. gondii* gene products are modified (Alonso et al., 2019). The functional relevance of myristoylation has been described for only a few *T. gondii* proteins, mainly in conjunction with adjacent palmitoylation that allows stable membrane attachment. These proteins include key signal mediators in parasite egress and invasion, e.g. CDPK3 (Garrison et al., 2012; McCoy et al., 2012), PKG (Brown et al., 2017), PKAr (Jia et al., 2017; Uboldi et al., 2018); proteins involved in parasite gliding, e.g. GAP45 and GAP70 (Frenal et al., 2010); division, e.g. ISP1, 2 and 3 (Beck et al., 2010); and correct rhoptry positioning required for invasion, e.g. ARO (Mueller et al., 2013). Collectively these studies show key roles for myristoylation throughout the parasite’s lytic cycle, but its function in the absence of palmitoylation or its relation to other P™s remains poorly described.

By combining metabolic tagging with orthogonal chemoproteomic tools and mass spectrometry (MS), we provide experimentally validated libraries of myristoylated as well as glycosylphosphatidylinositol (GPI) anchored proteins in *T. gondii*. We identify all the previously reported myristoylated proteins and novel substrates with heterogeneous localizations and variable functions across the lytic cycle. We validate the presence and elucidate the functional importance of myristoylation on two selected targets: the well characterized key signalling protein CDPK1 and the microneme protein MIC7. Utilizing conditional target depletion and complementation with wild-type (cWT) and myristoylation mutant (cMut) versions, we demonstrate that myristoylation is functionally important for CDPK1 and, somewhat surprisingly, MIC7, a predicted type-I-transmembrane protein. Taken together, our study points to unexpected and novel functions of myristoylation in *Toxoplasma* that extend beyond priming for palmitoylation and stable membrane attachment.

## Results

### Metabolic tagging allows for enrichment and visualization of myristoylated and GPI-anchored proteins in *T. gondii*

To visualize the extent of myristoylation in *T. gondii*, we adapted a metabolic tagging approach that has previously been applied to mammalian cells (Broncel et al., 2015; Thinon et al., 2014) and protozoan parasites (Wright et al., 2014; Wright et al., 2016; Wright et al., 2015). In this workflow, a myristic acid (Myr) analogue containing a terminal alkyne group (YnMyr) is added to cell culture upon infection with *Toxoplasma* (Figure 1A). The hydrophobic nature of YnMyr allows for cell membrane penetration, while the alkyne tag allows for NMT-mediated metabolic tagging of both host and parasite target proteins. Upon cell lysis, tagged proteins are liberated and conjugated to azide-bearing multifunctional capture reagents by a click reaction (Heal et al., 2011). The conjugation process introduces secondary labels, like biotin and fluorophores, allowing for target enrichment on streptavidin beads and visualization via in-gel fluorescence (igFL), respectively. To investigate the extent of YnMyr incorporation, intracellular tachyzoites were treated with either Myr or increasing concentrations of YnMyr for 16 h. Tagged proteins were conjugated to a capture reagent, resolved by SDS-PAGE, and visualized by igFL. Protein tagging *in vivo* was non-toxic and concentration-dependent without any detectable background (Figure S1A). In addition, it did not seem to depend on parasite localization inside or outside the host cell, and was efficiently out-competed by excess myristate, indicating that YnMyr works ‘on target’ (Figure S1B). To estimate the efficiency of target enrichment, we took advantage of the biotin moiety that enables a streptavidin-based pull down. We observed robust enrichment of protein targets in a YnMyr-dependent manner, and detected very little background in controls (Figure S1C).

**Figure 1.**
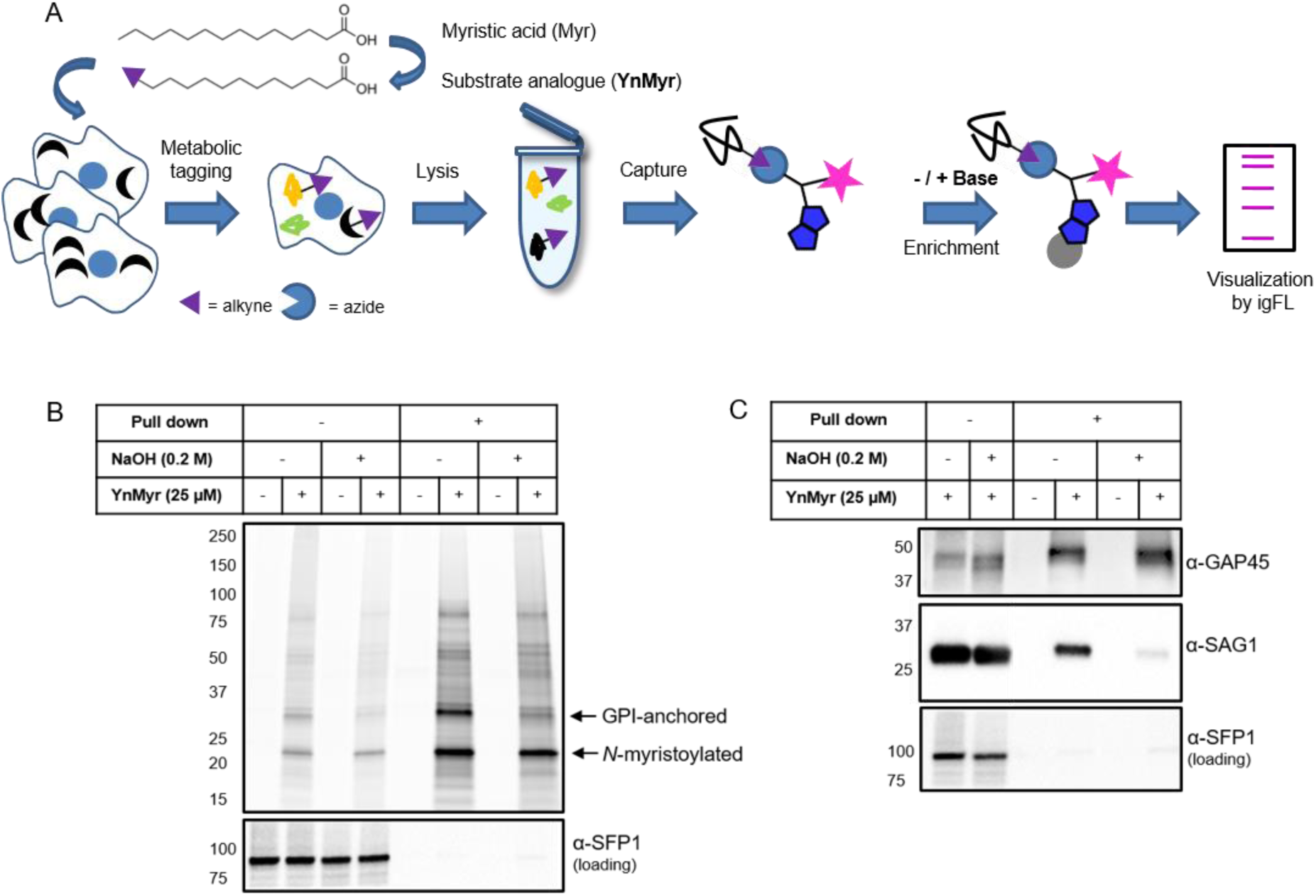
Metabolic tagging allows for enrichment and visualization of myristoylated and GPI-anchored proteins in *T. gondii.* (A) Metabolic tagging workflow. (B) In gel fluorescence visualization of YnMyr-dependent enrichment without and with the base treatment (top) and Western blot with α- SFP1 (TGGT1_289540) showing the loading control (bottom). (C) Western blot analysis of YnMyr-dependent pull down for known myristoylated and GPI-anchored proteins GAP45 and SAG1, respectively. See also Figure S1.

It has been reported that YnMyr can be incorporated not only at *N*-terminal glycines via amide bonds, but also through ester-linked incorporation of myristate into GPI anchors (Wright et al., 2014). These two distinct types of tagging can be readily distinguished by their different sensitivity to base treatment; amide bonds are stable in basic conditions, whereas ester bonds are hydrolysed. To visualize the extent of YnMyr incorporation into GPI anchors in *Toxoplasma*, we performed base treatment prior to enrichment of target proteins and observed a reduction of igFL signal for selected enriched bands (Figure 1B). To further validate the base treatment approach, we probed known *N*-myristoylated and GPI-anchored *Toxoplasma* proteins, GAP45 and SAG1, for their ability to be enriched in a base-dependent manner. In the absence of treatment, both proteins were robustly pulled down with YnMyr, while upon base treatment, only GAP45 remained enriched, confirming that it is a true myristoylation target (Figure 1C). Collectively, we confirmed that YnMyr is a robust and high-fidelity myristate analogue and demonstrated that it can be applied to profile both *N*-myristoylated and GPI-anchored proteins in live *Toxoplasma gondii.*

### Identification of the myristoylated proteome in *T. gondii*

To confidently identify myristoylated proteins in *Toxoplasma*, we applied state-of-the-art MS-based proteomics combined with validated chemical tools (Figure 2A; (Broncel et al., 2015; Speers and Cravatt, 2005; Thinon et al., 2014; Wright et al., 2014). We started with a small-scale pilot experiment to test our workflow, and differentiate between the myristoylation-based enrichment and the GPI-anchored targets. We metabolically labelled tachyzoites of the RH strain with either YnMyr or Myr each at 25 µM for 16 h. We then lysed the intracellular parasites and performed the click reaction with the azido biotin capture reagent **1** to facilitate YnMyr-dependent enrichment of tagged proteins. To distinguish myristoylated from GPI-anchored targets, we applied base treatment prior to the streptavidin-based pull down. Following trypsin digestion, we analysed samples by LC-MS/MS and performed label free quantification of enriched proteins. We quantified 2363 human and *Toxoplasma* proteins, 349 of which were parasite proteins with YnMyr intensities irrespective of base treatment (Table S1). To identify GPI-anchored proteins, we calculated log_2_ fold changes between base-treated and untreated samples (Figure S2A). To threshold we utilized the least extreme negative value (log_2_ fold change < −1) quantified from all Surface Antigen Proteins (SAGs) detected in our study, which are known to be GPI-anchored. This selection strategy yielded 52 targets, that included known and predicted GPI-anchored proteins (Table S1, Figure S2A). To identify myristoylated proteins we utilized a stringent selection method based on the robust YnMyr/Myr enrichment (log_2_ fold change > 2), the presence of a myristoylation motif (MG), and insensitivity towards base treatment. 56 proteins met these criteria, including those previously reported as myristoylated (Table S1). Analysis of supernatants from the pulled down samples did not reveal any substantial changes between the YnMyr and Myr proteomes, confirming that the observed target enrichment is not due to altered protein abundance (Figure S2B and Table S1).

**Figure 2.**
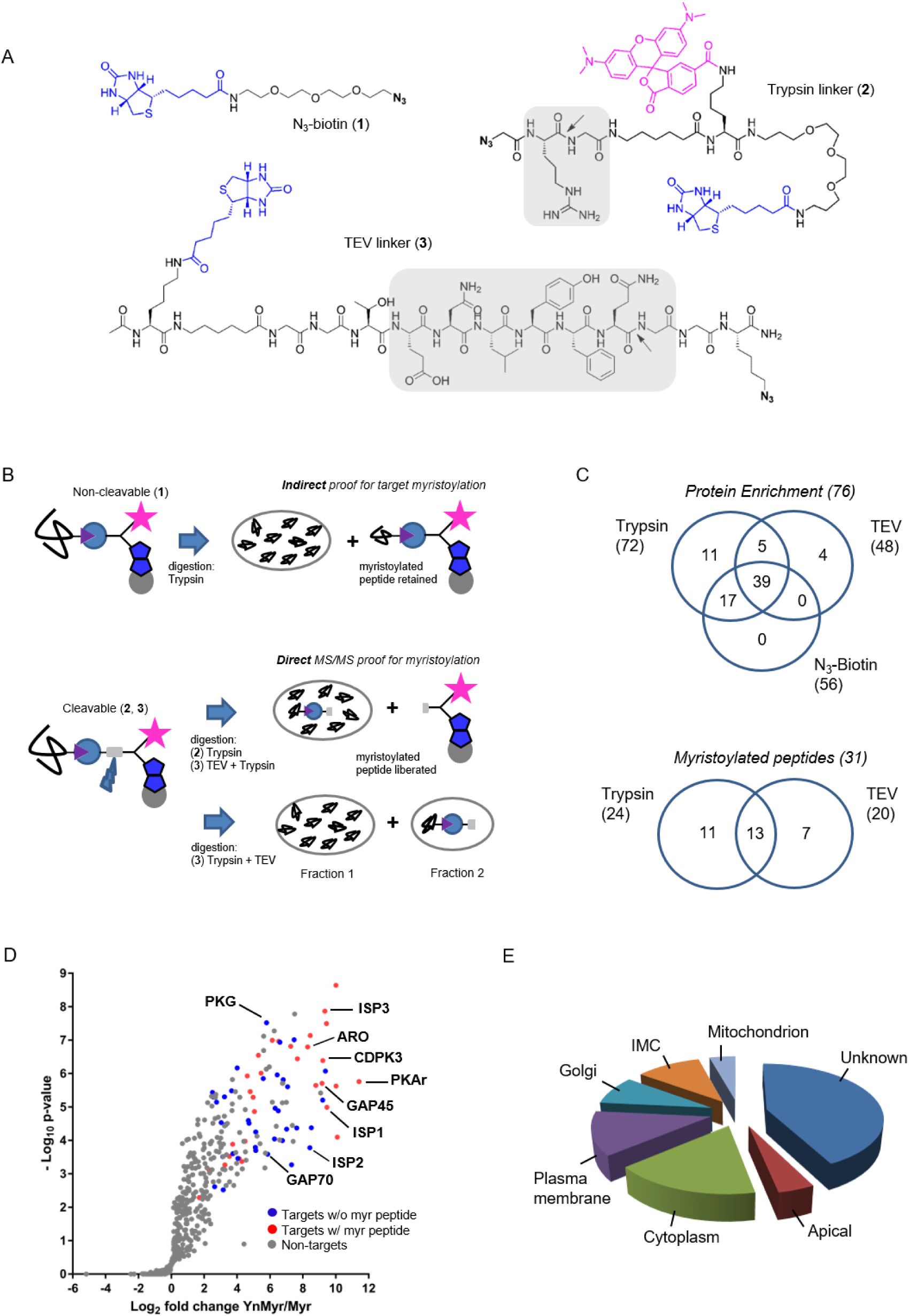
Identification of the myristoylated proteome in *T. gondii*. (A) Structures of capture reagents used in this study with key functional components highlighted: biotin and azide moieties in blue and bold, respectively, cleavable linkers in grey with the cleavage site indicated by arrows. (B) Schematic representation of the MS workflow using non-cleavable and cleavable capture reagents. (C) Top: Venn diagram illustrating the overlap between significantly YnMyr-enriched proteins identified with capture reagents used in this study. The number of significantly enriched proteins per reagent and in total is given in parenthesis. Bottom: Venn diagram showing the overlap in myristoylated peptide discovery between the two cleavable capture reagents used in this study. The number of modified peptides identified with each reagent and in total is given in parenthesis. (D) Label free quantification of the log_2_ fold changes in YnMyr enrichment over the Myr control plotted against the statistical significance for all parasite proteins detected in this study. Proteins with *N*-terminal glycine and significant, base-insensitive enrichment are highlighted in blue and red subject to the presence of a myristoylated peptide. All other identified proteins (background and GPI-anchors) are represented in grey. (E) Pie chart showing the distribution of reported and predicted (ToxoDB) cellular localization for the identified myristoylated proteins. See also Figure S2, Table S1, Table S2 and Table S3.

After successful testing of the metabolic tagging workflow, we performed a more elaborate MS experiment that utilizes cleavable capture reagents bearing either trypsin (reagent **2**) or TEV (reagent **3**) cleavable linkers (Figure 2A). In contrast to a non-cleavable reagent (e.g. reagent **1**) that provides only indirect proof of target myristoylation (Figure 2B), cleavable reagents allow for detection of myristoylated peptides in addition to peptides that originate from the enriched proteins (Figure 2B). This additional layer of confidence in target identification is especially important given the high level of non-myristoylation dependent background reported for metabolic tagging with YnMyr (Broncel et al., 2015; Wright et al., 2016; Wright et al., 2015). While reagent **2** has been validated as a tool for myristoylated protein and peptide discovery (Broncel et al., 2015), reagent **3** (Speers and Cravatt, 2005), which is expected to produce less background and improve myristoylated peptide discovery, has not previously been applied to study protein myristoylation. We therefore first tested reagent **3** in terms of YnMyr-dependent protein enrichment and observed robust pull down of potential targets (Figure S2C). We next generated samples for the MS workflow as described above but, instead of conjugating reagent **1**, we conjugated either **2** or **3**, each in biological triplicate, to tagged proteins via click reaction to enable myristoylation-dependent pull down. As depicted in Figure 2B, reagent **2** requires only a single step trypsin digestion to liberate both unmodified and myristoylated peptides in one pool. By contrast, reagent **3** requires both trypsin and TEV protease and, depending on the enzyme combination, releases unmodified and myristoylated peptides in either one (TEV I) or two (TEV II) separate fractions (Figure 2B). The TEV I strategy should address the common problem encountered with the on-bead trypsin digestion, i.e. the increased level of background from the non-specifically bound proteins, while the TEV II option should increase the modified peptide discovery. Following digestion, all samples were subjected to LC-MS/MS, and label free quantification was performed to identify proteins robustly enriched in YnMyr-dependent manner. This yielded 206 human and 117 *T. gondii* proteins bearing the MG myristoylation motif (Table S2). Within the parasite protein pool, we obtained statistically significant (FDR 1%, log_2_ fold change >2) enrichment in YnMyr over Myr controls for 72 targets using reagent **2** (Table S2). For reagent **3**, which was used in two different scenarios (TEV I and TEV II) resulting in larger variability between replicates, we utilized a fold change based threshold (log_2_ fold change > 2) and obtained 48 robustly enriched targets (Table S2). The overlap between reagents **2** and **3** reached 60% (Table S2), providing substantial confidence to the accuracy of our results. Reassuringly, we observed a ∼ 5 and ∼ 8-fold reduction in background in TEV I vs TEV II and TEV I vs reagent **2**, respectively, as shown by the number of proteins quantified in Myr controls (Table S2). Collectively we identified 76 significantly YnMyr-enriched proteins utilizing all three capture reagents (Figure 2C, Table S2).

We next focused on the identification of myristoylated peptides in samples processed with reagents **2** and **3**. Utilizing stringent criteria for the unbiased identification of the myristoylation adduct, as well as manual validation of the acquired MS/MS spectra, we identified 31 myristoylated peptides (Table S2), 24 of which were detected using reagent **2**, and 20 using reagent **3** (Figure 2C). None of these peptides were detected in Myr controls, and the myristoylation adduct was not identified on cysteines, confirming that YnMyr was not significantly incorporated into palmitoylation sites. Despite almost equal numbers of peptides detected by the two reagents, the overlap was only 40% (Figure 2C), confirming different specificities and the added value of orthogonal methods for modified peptide detection. As envisioned in our design strategy, we obtained a 30% increase in myristoylated peptide discovery in TEV II vs TEV I workflow (Figure S2D).

Finally, to generate a high confidence list of myristoylated proteins in *Toxoplasma*, we combined our results on both protein enrichment and the modified peptide levels. We filtered for proteins identified with at least two capture reagents or proteins for which we detected a lipid modified peptide. This resulted in 65 proteins, of which 48% have direct MS/MS evidence for protein myristoylation (Table S3, Figure 2D). Our target pool includes all proteins previously reported as myristoylated (Figure 2D) which indicates that this analysis covers a large fraction of the myristoylated proteome in *Toxoplasma*. The majority (90%) of our substrate pool represent novel targets of *Tg*NMT with important functions across the lytic cycle (Table S3), including CDPK1 (egress/invasion; (Lourido et al., 2010)); PPM5C (attachment; (Yang et al., 2019)); ARF1 and Rab5B (trafficking; (Kremer et al., 2013; Liendo et al., 2001)). We did not obtain any evidence for myristoylation on known secreted *Toxoplasma* proteins, such as rhoptry or dense granule proteins. Moreover, approximately one third of the reported targets are uncharacterized proteins, indicating that a large amount of unexplored myristoylation-related biology is still to be uncovered in *Toxoplasma*.

Identified targets showed heterogeneous localization (Figure 2E). We found proteins with known or predicted localization at the plasma membrane, as well as membrane-bound compartments (e.g. endoplasmic reticulum (ER) and Golgi apparatus). Stable attachment at membranes usually requires a double acylation, i.e. both myristoylation and palmitoylation (Wright et al., 2010). In agreement with this, 35% of our targets were previously reported to be palmitoylated ((Caballero et al., 2016; Foe et al., 2015); Table S3). Since palmitoylation is frequently enriched at the protein *N*-terminus, in close proximity to the myristate, we analysed the first 20 amino acid sequences of our targets (Figure S2E) and found that only about half of them had cysteines (sites of palmitoylation) and, hence, the potential for double acylation. This observation suggests that at least 50% of the target pool are likely only myristoylated at the *N*-terminus. Consistent with this, we found proteins with less defined localizations e.g. CDPK1, where myristoylation may serve a more discrete function beyond just a simple plasma membrane anchor.

We also compared our data with the myristoylated proteome of the related *P. falciparum* (Wright et al., 2014). We converted *Plasmodium* hits into *Toxoplasma* orthologues and compared the overlap of both species. This yielded 23 shared targets, which corresponds to 35% of the *Toxoplasma* and 62% of the *P. falciparum* experimentally validated myristoylated proteome (Figure S2F). Interestingly, 39 (60%) targets from the *Toxoplasma* dataset have orthologues in *P. falciparum* and 31 (48%) of them contain an MG motif, hinting at potentially unexplored *Pf*NMT substrates (Table S3). A further 26 (40%) of our targets did not have orthologues in *P. falciparum* (Table S3), indicating that they are unique to *Toxoplasma*, and therefore have the potential to uncover the parasite-specific biology.

### Myristoylation is important for CDPK1 function

CDPK1 is an essential regulator of microneme secretion and, therefore, a prerequisite for parasite motility, invasion and egress from host cells. Although the protein’s function has been thoroughly characterized (Lourido et al., 2013; Lourido et al., 2010; Ojo et al., 2010), it has not previously been shown as myristoylated. Accordingly, we first validated our finding by metabolic tagging and a myristoylation-dependent pull down after base treatment. As depicted in Figure 3A, CDPK1 was enriched in a base-independent manner which was further corroborated by the identification of the myristoylated peptide by MS (Figure 3B).

**Figure 3.**
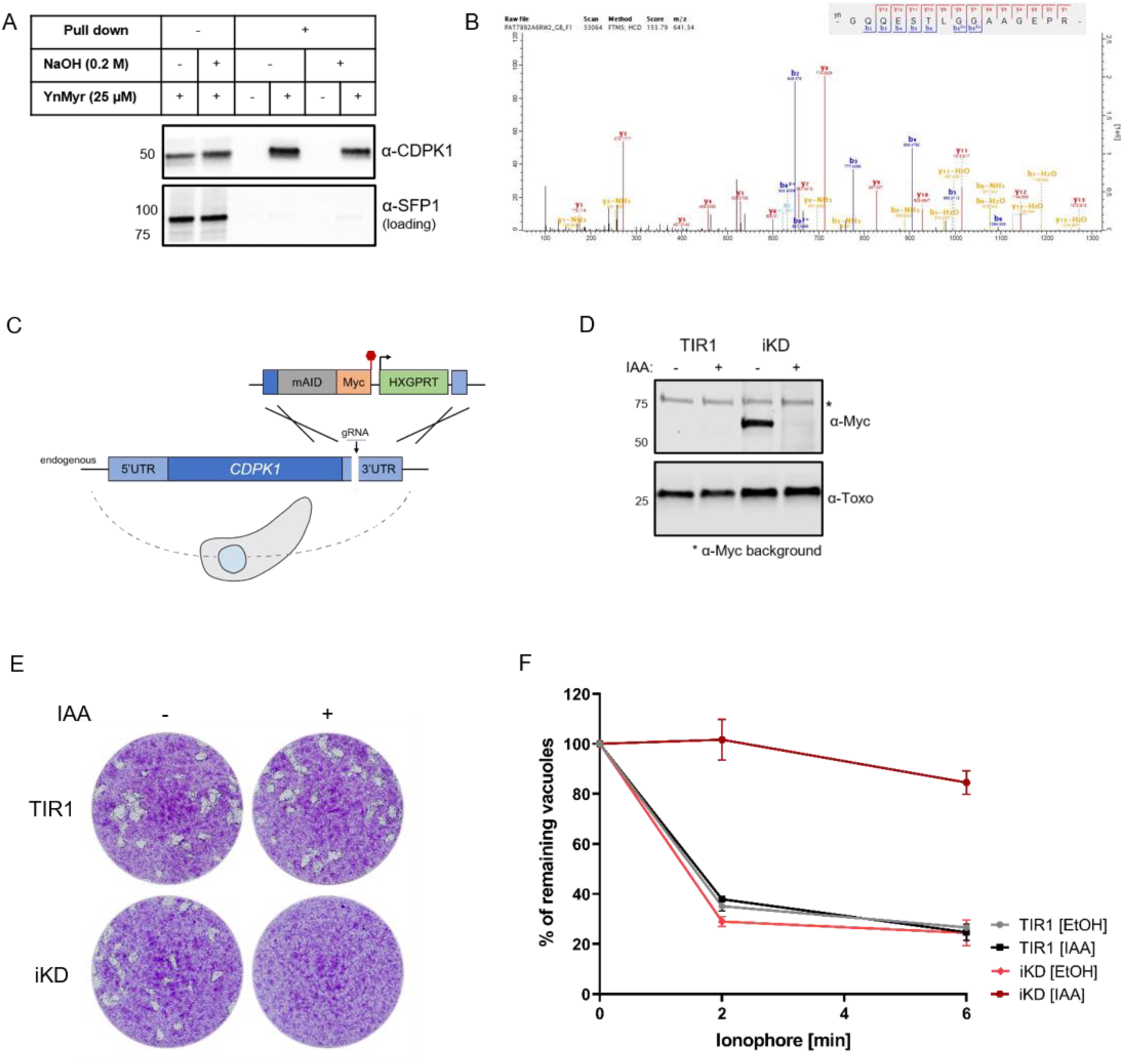
CDPK1 myristoylation and inducible knock-down. (A) Western blot analysis of YnMyr-dependent pull down for CDPK1 in base-treated and untreated samples. Base-insensitive pull down validates CDPK1 as a true myristoylation target. (B) MS/MS evidence for CDPK1 myristoylation. (C) Schematic representation of mAID-based knock-down strategy used for the conditional depletion of CDPK1. (D) Validation of IAA-dependent depletion of CDPK1 in the iKD line illustrated by Western blotting with α-Myc antibody. The band at 75 kD represents α-Myc-related background. α-*Toxoplasma* (Toxo) antibody was used as loading control. (E) Plaquing assays illustrating that CDPK1 is essential for the intracellular growth of *Toxoplasma.* Assay performed in three biological replicates, each in technical triplicate, representative images are shown. (F) Conditional depletion of CDPK1 abolishes ionophore-induced egress from host cells. Intracellular parasites were treated with IAA or vehicle (EtOH) for 2 h and egress was initiated by addition of 8 µM A23187. The number of intact vacuoles was monitored over the course of 6 min. Each data point is an average of two biological replicates, each in technical triplicate, error bars represent standard deviation. See also Figure S3.

To investigate the role of myristoylation in CDPK1 function, we generated an inducible knock-down (iKD) line using the mini auxin inducible degron (mAID) strategy (Brown et al., 2018), which allows for proteasomal degradation of mAID-tagged proteins in an indole acetic acid (IAA) dependent manner. To generate the CDPK1 iKD line, we *C*-terminally tagged endogenous CDPK1 in RHδku80 TIR1 parasites with a mAID-Myc cassette (Figure 3C). Successful mAID integration was verified by PCR (Figure S3), while tagged protein expression and IAA sensitivity was confirmed by immunoblot (Figure 3D). Introduction of the mAID cassette had no discernible fitness cost (Figure 3E), or impact on CDPK1’s function in egress (Figure 3F). By contrast, IAA-mediated depletion of CDPK1 had profound effects on the parasite lytic cycle (Figure 3E) and, as expected (Lourido et al., 2010), completely abolished ionophore-induced egress (Figure 3F). We complemented the iKD line by introducing HA-tagged WT (cWT) or myristoylation defective (cMut, G2>A) copies of *cdpk1* into the *uprt* locus (Figure 4A). We verified correct integration of both complementation constructs (Figure S4A), confirmed their equivalent and IAA-independent expression, as well as IAA sensitivity of the endogenous mAID-tagged CDPK1 within these lines (Figure 4B). In addition, we validated cWT and cMut lines biochemically by performing a YnMyr-dependent pull down and immunoblotting (Figure 4C). As expected, the endogenous copy of CDPK1 was efficiently enriched from both complemented lines, whereas the pull down of complements was only identified for the WT, but not the myristoylation mutant.

**Figure 4.**
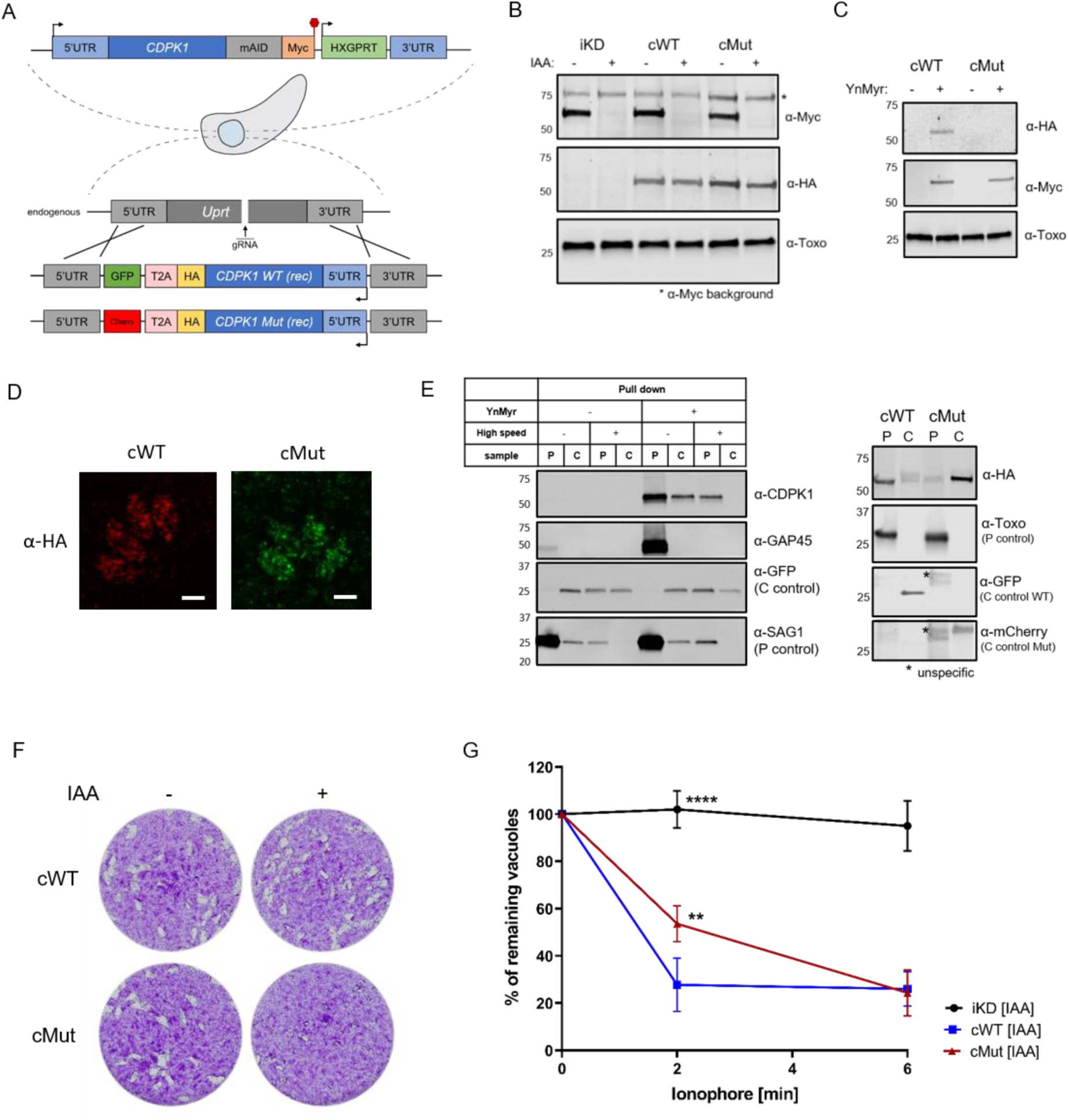
Myristoylation of CDPK1 is functionally important. (A) Complementation strategy used to evaluate the functional importance of CDPK1 myristoylation. Red hexagon represents STOP codon, rec – recodonized. (B) Western blot analysis demonstrating the IAA-dependent depletion of the endogenous copy of CDPK1 in the iKD, cWT and cMut lines (α-Myc) as well as equivalent and IAA-independent expression of the complements (α-HA). α-*Toxoplasma* (Toxo) antibody was used as loading control. (C) Biochemical validation of complemented lines by YnMyr-dependent pull down. Enrichment of WT and Mut copy of CDPK1 was evaluated by Western blotting with anti-HA antibody. The inducible copy of CDPK1 (anti-Myc) and Gra29 were used as enrichment and loading controls, respectively. (D) Localization of the complemented versions of CDPK1 within cWT and cMut by immunofluorescence analysis. Scale bar: 4 µm. (E) Myristoylation-dependent subcellular partitioning of CDPK1. (Left) Localization of YnMyr-enriched CDPK1 was evaluated using differential centrifugation. The partitioning into pellet [P] and cytosolic [C] fractions was revealed by Western blot (α-CDPK1) and compared to a doubly acylated GAP45. GFP and SAG1 were used as C and P controls, respectively. As only half of the cytosolic fraction was removed from the high speed pellet, the GFP signal is present in the latter. (Right) Complemented WT and Mut versions of CDPK1 were fractionated using exclusively high speed centrifugation and their partitioning between the P and C fractions revealed by Western blotting (α-HA). α-Toxoplasma (Toxo) antibody was used as P control whereas α-GFP and α-mCherry were used as C controls. (F) Plaquing assays demonstrating that myristoylation of CDPK1 is important in the intracellular growth of *Toxoplasma.* Assay performed in three biological replicates, each in technical triplicate, representative images are shown. (G) The lack of CDPK1 myristoylation delays ionophore-induced egress from host cells. Intracellular parasites were treated with IAA for 2 h and egress was initiated by addition of 8 µM A23187. The number of intact vacuoles was monitored over the course of 6 min. Each data point is an average of three biological replicates, each in technical triplicate, error bars represent standard deviation. Significance calculated using 1-way ANOVA with Tukey’s multiple comparison test, **p = 0.004, ****p < 0.0001. See also Figure S4.

Given that myristoylation is frequently reported to facilitate membrane association (Martin et al., 2011; Wright et al., 2010), we examined the localization of cWT and cMut CDPK1 by immunofluorescence analysis (Figure 4D). No clear differences were detected with a primarily punctate staining within the cytosol observed in both lines. As myristoylation-regulated changes in localization may be subtle, we explored the possible effects of myristoylation on CDPK1 localization using fractionation experiments (Figure 4E). First, we evaluated the partitioning pattern of the endogenous, myristoylated CDPK1. RHδku80 YFP expressing parasites were metabolically tagged with Myr or YnMyr and lysed in a hypotonic buffer to preserve intact membrane structures. Next, lysates were pelleted by centrifugation (16,000 × g) and the resulting cytosolic fraction was subjected to an additional high speed (100,000 × centrifugation step. Each fraction was then clicked to a capture reagent, pulled down and myristoylation-dependent partitioning revealed by Western blotting. In contrast to the doubly acylated GAP45, which was present exclusively in the pellet, CDPK1 was observed in both the pellet and cytosolic fractions of the first spin (Figure 4E, left). Following further centrifugation of the cytosolic fraction, CDPK1 partitioned exclusively into the pellet (Figure 4E, left), suggesting potential association with some vesicular structures or higher molecular weight complexes. We next used both cWT and cMut lines to elucidate any myristoylation dependent changes to CDPK1 localization. While cWT could be found exclusively in the 100,000 × g pellet, removal of myristoylation in the cMut caused a release of the pellet associated protein into cytosolic fraction (Figure 4E, right).

To evaluate the role of CDPK1 myristoylation in parasite fitness, we performed plaque assays (Figure 4F). In the presence of the endogenous copy of CDPK1, both complemented lines developed normally. Upon IAA-mediated depletion of endogenous CDPK1, however, we observed a substantial decrease in cMut plaque size. This finding demonstrates that one or more steps of the *Toxoplasma* lytic cycle are negatively affected by a loss of CDPK1 myristoylation. In light of CDPK1’s known function in egress, we next explored whether CDPK1 myristoylation might regulate the parasite’s ability to leave its host cell. In the absence of auxin, cWT, cMut, and the phenotypic control line (iKD), egressed within 2 min of ionophore treatment (Figure S4B). While the cWT line maintained similar egress kinetics upon addition of IAA, the cMut line showed a significant delay after 2 min of treatment (Figure 4G). This egress delay was overcome at the 6 min time point (Figure 4G), suggesting that CDPK1 myristoylation is important for ionophore-induced egress, but not essential. Collectively, our results demonstrate an important role for CDPK1 myristoylation in the lytic cycle, which may be in part mediated by a defect in egress.

### MIC7 is myristoylated and is important for *T. gondii* lytic cycle

Within our high confidence target pool, we also identified the microneme protein MIC7. MIC7 has been reported to be a putative type-I-transmembrane protein, comprising an *N*-terminal signal peptide, five EGF-like domains, a membrane-spanning region, and a short cytoplasmic tail (Meissner et al., 2002). As MIC signal peptides are typically co-translationally cleaved upon entry into the ER (Soldati et al., 2001), the presence of a myristate within the classical signal sequence of MIC7 was unusual. In addition, MIC7 has been shown to be predominantly expressed in bradyzoites (Meissner et al., 2002), the lifecycle stage responsible for the chronic phase of *T. gondii* infection. As our experiments were performed exclusively in tachyzoites, the stage responsible for acute infection, the presence of MIC7 within our dataset could represent a potential false positive identification. To address this inconsistency, we performed MS-based quantification of bradyzoite and tachyzoite proteomes (unpublished dataset). The log_2_ fold changes in protein abundance for MIC7 and the bradyzoite-specific marker, MAG1 ((Tu et al., 2019); Figure 5A, Table S4) revealed that in contrast to MAG1, MIC7 is expressed in tachyzoites, supporting the MS and transcriptional evidence in ToxoDB. We next aimed to validate the protein’s myristoylation as our MS screen did not identify a myristoylated peptide for MIC7. We ectopically expressed HA-tagged WT and myristoylation mutant (Mut, G2G3>KA) copies of MIC7 under control of either the endogenous or tubulin promoter, metabolically labelled cultures with YnMyr and performed a myristoylation-dependent pull down on lysates. Only WT but not the Mut was enriched in this manner (Figure 5B), showing that MIC7 is indeed myristoylated.

**Figure 5.**
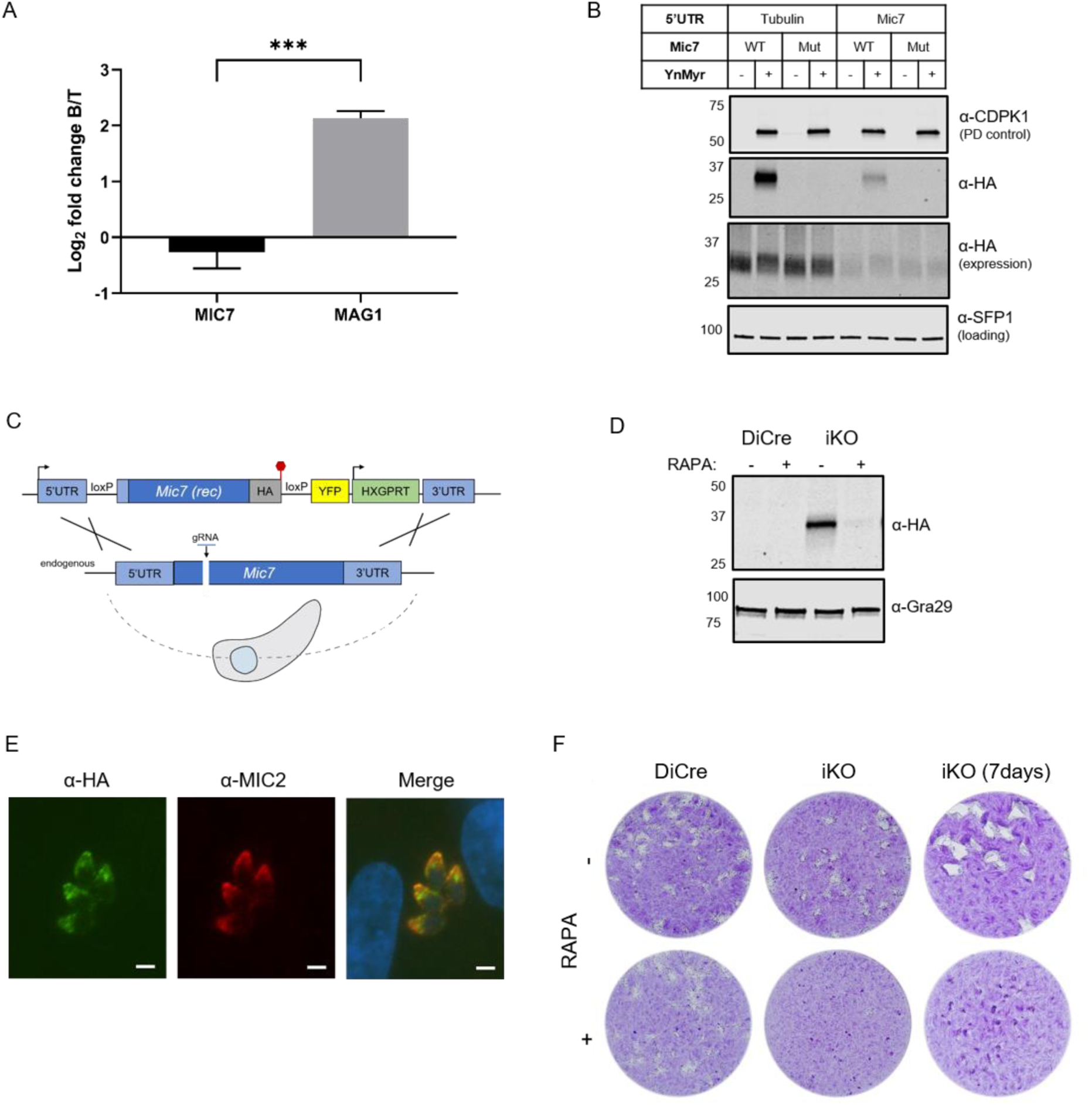
MIC7 is myristoylated and is important for *T. gondii* lytic cycle. (A) MS-based quantification of MIC7 and MAG1 abundance in tachyzoites [T] and bradyzoites [B] of *T. gondii*. Significance calculated using two-tailed Student’s t-test, ***p = 0.0002, N=3. (B) MIC7 is myristoylated as shown by YnMyr-dependent pull down and Western blotting with α-HA antibody. CDPK1 and SFP1 (TGGT1_289540) are used as enrichment and loading controls, respectively. (C) Schematic representation of the DiCre/loxP-based iKO strategy used for the conditional depletion of *mic7*. Red hexagon represents STOP codon, rec - recodonized. (D) Validation of RAPA-dependent depletion of MIC7 in the iKO line illustrated by Western blotting with α-HA antibody. Gra29 was used as loading control. (E) Co-localization of the inducible version of MIC7 (green) with the micronemal marker MIC2 (red) in the iKO line by immunofluorescence analysis. Scale bar: 3 µm. (F) Plaquing assays illustrating that MIC7 is important, but not essential for the intracellular growth of *Toxoplasma.* Assay was performed for 5 days (where not indicated) in three biological replicates, each in technical triplicate, representative images are shown. See also Figure S5.

To investigate the functional relevance of MIC7 and its myristoylation, we created an inducible knock-out (iKO) line using the DiCre/loxP system (Andenmatten et al., 2013) that we recently optimised in RHδku80 parasites (Hunt, 2019). The *mic7* coding sequence was replaced with a floxed, HA-tagged copy of the gene that could be excised upon rapamycin (RAPA) treatment (Figure 5C). We verified correct integration at the endogenous locus (Figure S5A) and confirmed RAPA-induced excision (Figure S5B) by PCR. At the protein level, MIC7 was efficiently depleted 24 h post RAPA treatment (Figure 5D). Correct trafficking of MIC7 to micronemes was verified by the co-localization with the micronemal marker MIC2 (Figure 5E). Upon deletion of *mic7*, parasites no longer formed detectable plaques in host cell monolayers after 5 days in culture, but we could observe very small plaques emerging after 7 days (Figure 5F). Collectively these results demonstrate an important, but non-essential, role for MIC7 in the lytic cycle.

### Myristoylation of MIC7 plays a role in the invasion of host cells

To investigate where in the lytic cycle MIC7 plays a role, and test the functional relevance of *N*-terminal myristoylation, we introduced cWT and cMut copies of *mic7* into the *uprt* locus of the iKO line (Figure 6A). Inserts coding for Ty1-tagged cWT or cMut MIC7 were correctly integrated (Figure S6A) and both complemented lines retained efficient RAPA-induced *mic7* excision (Figure S6B) and depletion of the endogenous protein (Figure 6B). After confirming equivalent and RAPA-insensitive expression of cWT and cMut (Figure 6B), we validated both lines in terms of their myristoylation-dependent enrichment and showed that only the cWT MIC7 was selectively pulled down after metabolic tagging with YnMyr (Figure 6C).

**Figure 6.**
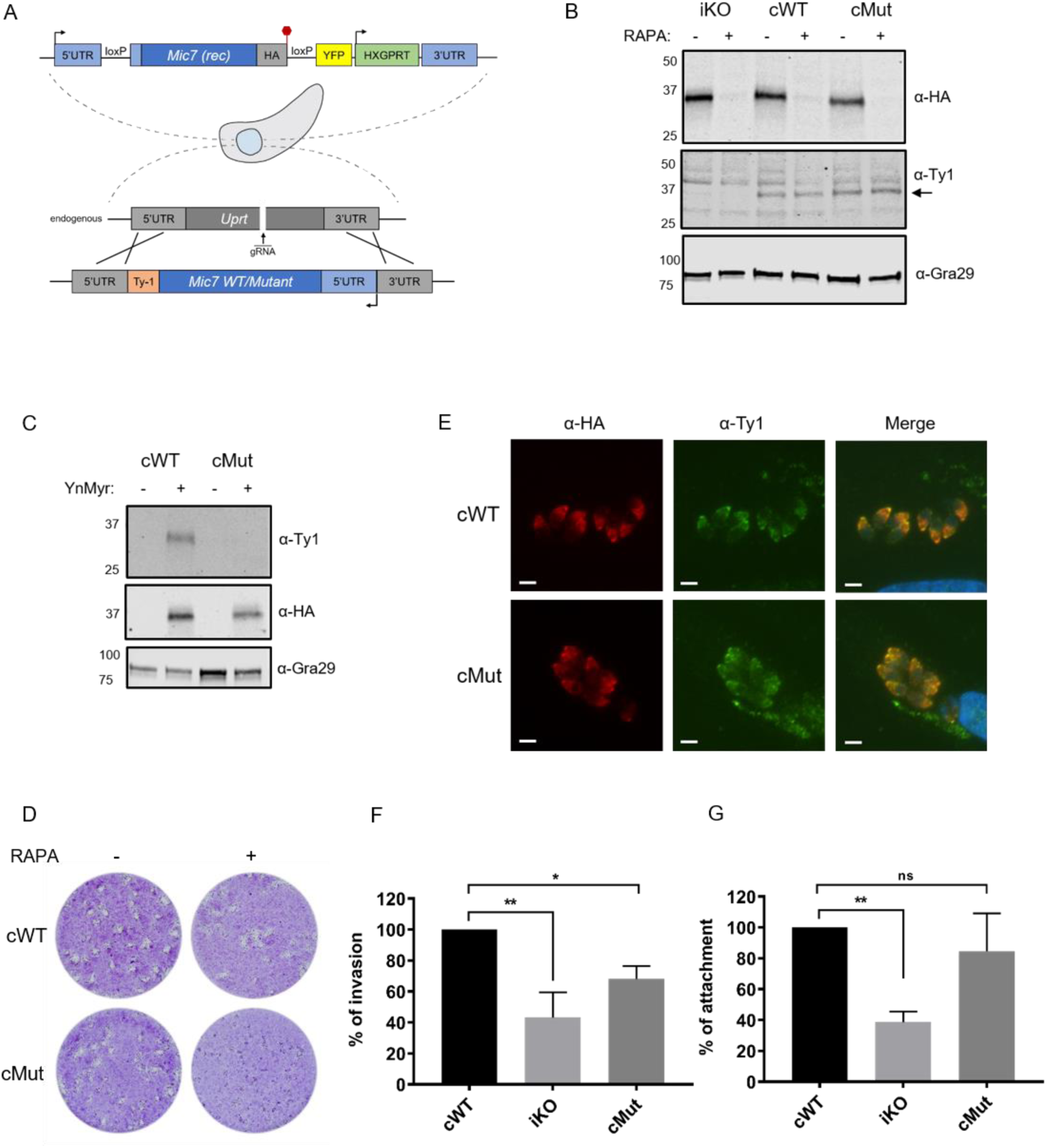
Myristoylation of MIC7 plays a role in the invasion of host cells. (A) Complementation strategy used to evaluate the functional importance of MIC7 myristoylation. Red hexagon represents STOP codon, rec – recodonized. (B) Western blot analysis demonstrating the RAPA-dependent depletion of the endogenous copy of MIC7 in the iKO, cWT and cMut lines (α-HA) as well as equivalent and RAPA-independent expression of the complements (α-Ty1). Gra29 was used as loading control. (C) Biochemical validation of complemented lines by YnMyr-dependent pull down. Enrichment of WT and Mut copy of MIC7 was evaluated by Western blotting with α-Ty1 antibody. The inducible copy of MIC7 (α-HA) and Gra29 were used as enrichment and loading controls, respectively. (D) Plaquing assay demonstrating that myristoylation of MIC7 is important in the intracellular growth of *Toxoplasma.* Assay performed for 5 days in three biological replicates, each in technical triplicate, representative images are shown. (E) Co-localization of the inducible version of MIC7 (red) with the complemented WT and Mut (green) by immunofluorescence analysis. Scale bar: 5 µm. (F) Quantification of invasion efficiency in the RAPA-treated cWT, iKO and cMut lines. Figure shows the average of three biological replicates, each in technical duplicate, error bars represent standard deviation. Significance calculated using 1-way ANOVA with Dunnett’s multiple comparison test, **p = 0.001, *p = 0.018. (G) Quantification of attachment efficiency in the RAPA-treated cWT, iKO and cMut lines. Figure shows the average of three biological replicates, each in technical duplicate, error bars represent standard deviation. Significance calculated using 1-way ANOVA with Dunnett’s multiple comparison test, **p = 0.004, ns = not significant. See also Figure S6.

We next evaluated the role of MIC7 myristoylation in the parasite lytic cycle. While complementation with WT MIC7 rescued the iKO phenotype upon RAPA treatment, cMut parasites formed substantially smaller plaques under equivalent conditions (Figure 6D). This demonstrates that myristoylation plays a key role in regulating MIC7 function. To shed further light on the nature of this myristoylation-dependent phenotype, we investigated the co-localization of endogenous MIC7 with Ty1-tagged cWT and cMut (Figure 6E). Both complementation isoforms localized to the micronemes, indicating that the myristate is not required for the trafficking of MIC7 to this organelle.

Given the well-established role of microneme proteins in facilitating host cell penetration we explored whether myristoylation of MIC7 may be important for invasion. We treated iKO, cWT and cMut parasites with RAPA and performed a red/green assay (Huynh et al., 2003) which can distinguish invaded from attached parasites. As shown in Figure 6F, we observed efficient invasion of host cells by the cWT parasites. This was not the case in the iKO and cMut lines, where invasion was reduced by 57% and 32%, respectively. Compared to the cWT line, we also observed a consistent 61% drop in the total number of iKO parasites, which suggests a defect in the attachment to host cells. A modest but non-significant reduction of 15% in attachment was observed in the cMut strain. Collectively, these results indicate that MIC7 plays an important role in *Toxoplasma* propagation by facilitating parasite attachment and subsequent entry into host cells. Furthermore, myristoylation is not required for sorting MIC7 to the micronemes but appears to be important for the protein’s function in invasion of host cells.

## Discussion

Our understanding of myristoylation and its functional consequences in *Toxoplasma* is hampered by the limited knowledge of NMT substrates. Utilizing an MS-approach which allows for increased target coverage and provides direct evidence for target modification, we describe here the first experimentally validated myristoylated proteome in *T. gondii*. Despite the complex nature of our samples, consisting of both human and parasite proteins, our discovery includes all proteins previously reported to be myristoylated in *Toxoplasma* as well as novel *Tg*NMT substrates. The fact that these proteins are functionally diverse, and involved in all steps of the lytic cycle highlights the importance of myristoylation in *Toxoplasma* biology. In light of this, we predict that NMT inhibition would exert severe pleiotropic effects on the *Toxoplasma* lytic cycle. Although potent and selective *Tg*NMT inhibitors are yet to be reported, extensive work in other protozoan parasites (Schlott et al., 2019; Wright et al., 2014; Wright et al., 2016; Wright et al., 2015) demonstrates that selective NMT inhibition could provide an attractive strategy to combat infection.

Despite the frequent coincidence of myristoylation with palmitoylation, our target pool overlapped only moderately with published palmitoylome datasets (Caballero et al., 2016; Foe et al., 2015). A lack of cysteine enrichment within the 20 *N*-terminal residues of half our targets indicates they may be myristoylated only, suggesting that their myristoylation can serve more discrete functions than just a priming site for the palmitate. Such alternate functions could include reversible membrane binding by the conformation regulated exposure of the myristate as shown for mammalian ARF1 (Goldberg, 1998) or involvement in PPIs, as demonstrated for the picornavirus capsid assembly (Chow et al., 1987). To further investigate the importance of myristoylation in *Toxoplasma* and validate our study, we selected two targets, CDPK1 and MIC7, for analysis.

We have shown that CDPK1 is myristoylated, and that this modification subtly regulates its localization. Contrary to previous reports, which described the protein as cytosolic or even nuclear (Ojo et al., 2010; Pomel et al., 2008), our findings indicate that myristoylated CDPK1 is associated with structures that can be fractionated from the cytoplasmic protein pool by differential centrifugation. We predict that these are membranous structures, visualised as puncta within the cytoplasm, but cannot exclude the possibility that they are protein complexes. Irrespective of their nature, their association with CDPK1 is lost upon removal of myristoylation. This phenomenon could have major impact on the CDPK1’s ability to access its targets and it would be interesting to test in the future whether there are subsets of CDPK1 substrates that depend on its myristoylation state. The elegant use of engineered parasites to identify direct targets of CDPK1 (Lourido et al., 2013) may allow such analysis.

CDPK1 has been shown to be essential for *Toxoplasma* fitness, including egress (Lourido et al., 2010). The importance of CDPK1 in these processes was also confirmed here. Mutating the myristoylation site, however, had only a modest effect in ionophore-induced egress assays, while in plaque assays this modification appears very important. It is possible that myristoylation of CDPK1 could be required for processes that are not egress related, e.g. host cell invasion. In line with that, a function of CDPK1 to control the actomyosin system and extrusion of the conoid was recently suggested (Tosetti et al., 2019). Alternatively, the forced nature of induced egress could lead to compensatory effects, potentially by other kinases. The plausible candidates could be CDPK3 (Treeck et al., 2014), PKG (Brown et al., 2017), or CDPK2a, which we found robustly YnMyr-enriched in all our experiments, but since it lacks the *N*-terminal MG motif, it was excluded from our high confidence dataset.

Microneme proteins are key factors in *Toxoplasma* propagation, as they are involved in parasite egress, motility and host cell invasion (Soldati et al., 2001). Here we show that MIC7 is indeed an important protein required for completion of the lytic cycle. How MIC7 functions in this process requires further work, but our data indicate that attachment and invasion of host cells are severely impacted upon its deletion. As is the case for all known microneme proteins, also MIC7 was reported to possess a signal peptide (Meissner et al., 2002). Here we show that the presence of a signal sequence within MIC7 is unlikely, instead the protein is *N*-terminally myristoylated. The fact that myristoylation is not required for sorting MIC7 to the micronemes and appears to be important in invasion of host cells opens up many questions regarding the precise function of this modification at the MIC7 *N*-terminus. While it is known that proteins can be sorted to secretory pathway by virtue of recessed signal or leader peptides, this has not been reported for microneme proteins. If the myristate is not important for trafficking, it could facilitate PPIs with other microneme proteins. It is also plausible that the myristate could contribute to invasion through direct insertion into the host cell membrane. While the experimental evidence for these hypotheses is still pending, it has been shown that certain viruses can use myristoylation to promote PPIs required for capsid assembly (Simons et al., 1993) as well as to enter host cells (Maurer-Stroh and Eisenhaber, 2004). Interestingly, blast analysis of the MIC7 sequence against the *Toxoplasma* proteome identifies a paralogue in bradyzoites (TGME49_315520). This protein also contains an MG motif, a predicted transmembrane domain with a short cytoplasmic tail and no predicted signal peptide, hinting at the presence of a stage-specific MIC7-like invasion ligand, as has been observed for AMA1 (Lamarque et al., 2014).

In conclusion, we have performed the first high throughput screening of protein myristoylation in *Toxoplasma gondii* providing a useful resource of experimentally validated myristoylated as well as GPI-anchored proteins. Furthermore, we have shown the functional relevance of myristoylation, that is unrelated to priming for palmitoylation, on two proteins important in *Toxoplasma* lytic cycle. This demonstrates how our discovery can serve as a tool in target-specific investigations that can ultimately help to unravel the exciting biology of the parasite.

## Acknowledgments

We would like to thank the Tate, Lourido, Carruthers, and Sibley labs for sharing reagents. We also thank members of the following Science Technology platforms at The Francis Crick Institute: Proteomics, Peptide Synthesis, and High Throughput Screening. This work was supported by funding from The Francis Crick Institute (https://www.crick.ac.uk/), which receives its core funding from Cancer Research UK (FC001189; FC001999), the UK Medical Research Council (FC001189; FC001999) and the Wellcome Trust (FC001189; FC001999) as well as the National Institute of Health grant (NIH-R01AI123457).

## Author contributions

Conceptualization, M.B. and M.T; Methodology, M.B., C.D., and M.T.; Validation, M.B.; Investigation, M.B., C.D., A.H., B.W., and J.Y.; Resources, S.F.; Data curation, M.B.; Writing – original draft, M.B. and M.T.; Writing – Review and Editing, M.B., C.D., J.Y., and M.T.; Visualization, M.B., A.H., and J.Y.; Supervision, M.T.; Funding acquisition, M.T.

## Declaration of Interests

The authors declare no competing interests

## Materials and Methods

### General

Reagents: CuSO_4_, TCEP, TBTA, buffer salts, DTT, iodoacetamide, DMSO, BSA, Triton-X100 and Tween-20 were from Sigma Aldrich. Azide-PEG3-biotin was from Sigma Aldrich. Peptide synthesis coupling reagents HATU and HCTU were from Fluorochem and Merck, respectively. MS-grade water, acetonitrile, methanol, TFA and formic acid were from Thermo Scientific.

### Plasmid generation

Primers used throughout this study are listed in Table S5. Plasmid sequences were confirmed by Sanger sequencing (Eurofins Genomics).

To generate the *Mic7* iKO plasmid, pG140_MIC7_HA_iKO_loxP100, the *Mic7* 5’UTR with a loxP site inserted 100 bp upstream of the *Mic7* start codon, and a recodonised *Mic7* cDNA-HA sequence, were synthesized (GeneArt strings, Life Technologies). These DNA fragments were Gibson cloned into the ApaI/PacI digested parental vector p5RT70loxPKillerRedloxPYFP-HX (Andenmatten et al., 2013) to generate an intermediate plasmid. The *Mic7* 3’UTR was subsequently amplified from genomic DNA using primers 1 and 2, while *mCherry* flanked by *Gra* gene UTRs was amplified from pTKO2C (Caffaro et al., 2013) using primer pair 3/4. The resulting fragments were Gibson cloned into the SacI-digested intermediate plasmid to generate pG140_MIC7_HA_iKO_loxP100.

To generate the complementation construct pUPRT_MIC7_Ty1, the *Mic7* sequence flanked by its 5’UTR was amplified from genomic DNA using primer pair 5/6. In parallel, the *Uprt* targeting vector pUPRT_HA (Reese et al., 2011) was amplified by inverse PCR using primers 7 and 8. The resulting PCR amplicons were Gibson cloned to generate pUPRT_MIC7_Ty1. Primers 5 and 8 comprise overhangs to facilitate introduction of a Ty1 tag 3’ of the *Mic7* sequence.

To generate the complementation construct pUPRT_MIC7(G2K/G3A)_Ty1, the *Mic7* 5’UTR and *Mic7* endogenous sequence were amplified using primer pairs 9/10 and 5/11, respectively. In parallel, the *Uprt* targeting vector pUPRT_HA (Reese et al., 2011) was amplified by inverse PCR using primers 7 and 8. The resulting PCR amplicons were Gibson cloned to generate pUPRT_MIC7(G2K/G3A) _Ty1. Primers 9 and 11 comprise overhangs that introduce point mutations G2K and G3A, while primers 5 and 8 introduce a Ty1 tag 3’ of the *Mic7* sequence.

To generate pSag1_Cas9-U6_sgMIC7, the pSag1_Cas9-U6_sgUPRT ((Shen et al., 2014); Addgene plasmid # 54467) vector was amplified by inverse PCR using primers 12 and 13. Primer 13 comprises a sequence extension that replaces the *Uprt*-targeting sgRNA with a sgRNA sequence targeting *Mic7*. The resulting linear fragment was circularized using KLD reaction buffer (NEB) as per manufacturer’s instructions.

To generate pGra_5’UTRMIC7_MIC7_HA, the 5’UTR of *Mic7* was amplified from gDNA using primer pair 53/54, and recodonised *Mic7* sequence was amplified from pG140_MIC7_HA_iKO_loxP100 using primers 55 and 56. In parallel, the vector pGra_ApiAT5-3_HA (Wallbank et al., 2019) was amplified by inverse PCR using primer pair 57/58. The 3 resulting PCR amplicons were Gibson assembled to generate pGra_5’UTRMIC7_MIC7_HA.

To generate pGra_5’UTRMIC7_MIC7(G2K/G3A) _HA, the 5’UTR of *Mic7* was amplified from gDNA using primer pair 53/54, and recodonised *Mic7(G2K/G3A)* sequence was amplified from pG140_MIC7_HA_iKO_loxP100 using primers 59 and 56. Primer 59 was used to introduce the point mutations G2K and G3A into the *Mic7* recodonised sequence. In parallel, the vector pGra_ApiAT5-3_HA (Wallbank et al., 2019) was amplified by inverse PCR using primer pair 57/58. The 3 resulting PCR amplicons were Gibson assembled to generate pGra_5’UTRMIC7_MIC7(G2K/G3A)_HA.

To generate pGra_5’UTRTUB_MIC7 _HA, the *Tub* 5’UTR was amplified from gDNA using primer pair 60/61, and recodonised *Mic7* sequence was amplified from pG140_MIC7_HA_iKO_loxP100 using primers 62 and 56. In parallel, the vector pGra_ApiAT5-3_HA (Wallbank et al., 2019) sequence was amplified by inverse PCR using primer pair 57/63. The 3 resulting PCR amplicons were Gibson assembled to generate pGra_5’UTRTUB_MIC7 _HA.

To generate pGra_5’UTRTUB_MIC7(G2K/G3A)_HA, the *Tub* 5’UTR was amplified from gDNA using primer pair 60/61, and recodonised *Mic7(G2K/G3A)* sequence was amplified from pG140_MIC7_HA_iKO_loxP100 using primers 64 and 56. Primer 64 was used to introduce the point mutations G2K and G3A into the *Mic7* recodonised sequence. In parallel, the vector pGra_ApiAT5-3_HA (Wallbank et al., 2019) was amplified by inverse PCR using primer pair 57/63. The 3 resulting PCR amplicons were Gibson assembled to generate pGra_5’UTRTUB_MIC7(G2K/G3A) _HA.

To generate pTUB1_YFP_mAID_Myc, the pTUB1_YFP_mAID_3HA ((Brown et al., 2017); Addgene plasmid #87259) vector was amplified by inverse PCR using primer pair 14/15 to substitute the 3HA tag sequence for a Myc tag encoding sequence. The resulting linear fragment was circularized using KLD reaction buffer (NEB) as per manufacturer’s instructions.

To generate pSag1_Cas9-U6_sg3’UTRCDPK1, the pSag1_Cas9-U6_sgUPRT ((Shen et al., 2014); Addgene plasmid # 54467) vector was amplified by inverse PCR using primers 12 and 16. Primer 16 comprises a sequence extension that replaces the *Uprt*-targeting sgRNA with a sgRNA sequence targeting the 3’UTR of *CDPK1*. The resulting linear fragment was circularized using KLD reaction buffer (NEB) as per manufacturer’s instructions.

To generate the complementation construct pUPRT_CDPK1_ HA_T2A_GFP, the *CDPK1* 5’UTR was amplified from genomic DNA using primer pair 17/18. In parallel, recodonised *CDPK1* cDNA-HA sequence was synthesised (GeneArt strings, Life Technologies) and amplified with appropriate overhangs using primers 19 and 20. Sequence encoding T2A-GFP was amplified from an in-house unpublished plasmid using primer pair 21/22. The three resulting fragments were Gibson cloned into the PacI-linearised pUPRT_HA (Reese et al., 2011) vector.

To generate the complementation construct pUPRT_CDPK1(G2A)_HA_T2A_mCherry, *CDPK1* 5’UTR was amplified from genomic DNA using primer pair 23/18. Recodonised *CDPK1-HA* was amplified from pUPRT_CDPK1_ HA_T2A_GFP with appropriate overhangs using primers 24 and 25. Primers 23 and 25 were used to introduce a G2A point mutation within *CDPK1*. Sequence encoding *T2A-mCherry* was amplified from an in-house unpublished plasmid using primer pair 21/26. The three resulting PCR amplicons were Gibson cloned into the PacI-linearized pUPRT_HA (Reese et al., 2011) vector.

### Parasite strain generation

Freshly harvested parasites were transfected by electroporation (1500 V) using the Gene Pulser Xcell system (Bio-Rad) as previously described (Soldati and Boothroyd, 1993).

To generate the inducible MIC7 knock-out strain (RH DiCreΔku80Δhxgprt_loxPMIC7_HA, referred to here as iKO MIC7), the plasmid pG140_MIC7_HA_iKO_loxP100 was linearized using PciI and co-transfected with pSag1_Cas9-U6_sgMIC7 into the RH DiCreΔku80Δhxgprt strain (Andenmatten et al., 2013). Recombinant parasites were selected 24 h post transfection by addition of mycophenolic acid (MPA; 25µg/mL) and xanthine (XAN; 50 µg/mL) to culture medium. Resistant non-fluorescent parasites were cloned, and successful 5’ and 3’ integration at the *Mic7* locus was confirmed using primer pairs 27/28 and 29/30, respectively. Absence of the endogenous *Mic7* locus was confirmed using primers 43 and 44. Rapamycin-induced excision of the *loxPMic7* sequence was confirmed using primer pair 45/46.

To complement the iKO strain with MIC7-expressing constructs, pUPRT_MIC7_Ty1 and pUPRT_MIC7(G2K/G3A)_Ty1 plasmids were linearized with ScaI and individually co-transfected with pSAG1_Cas9-U6_sgUPRT. Transgenic parasites were subjected to 5’-fluo-2’-deoxyuridine (FUDR) selection (5 µM) 24 h post transfection. Resistant parasites were cloned, and successful 5’ and 3’ integration was confirmed using primer pairs 31/32 and 33/34. Disruption of the endogenous *Uprt* locus was confirmed using primer pair 47/48.

To generate lines that express WT and myristoylation mutant (G2K/G3A) MIC7 ectopically, plasmids pGra_5’UTRMIC7_MIC7_HA, pGra_5’UTRMIC7_MIC7(G2K/G3A)_HA, pGra_5’UTRTUB_MIC7_HA, and pGra_5’UTRTUB_MIC7(G2K/G3A)_HA were linearized using NotI and individually transfected into the RH Δhxgprt strain. Recombinant parasites were selected 24 h post transfection by addition of mycophenolic acid (MPA; 25µg/mL) and xanthine (XAN; 50 µg/mL) to culture medium.

To generate the inducible CDPK1 knock-down strain (RH TIR-1-3FLAG_CDPK1_mAID_Myc, referred to here as iKD CDPK1), the sequence encoding *mAID_Myc_HXGPRT* was PCR amplified from pTUB1_YFP_mAID_Myc using primer pair 35/36, and co-transfected into the RH TIR1-3FLAG (Brown et al., 2018) strain with pSag1_Cas9-U6_sg3’UTRCDPK1. Recombinant parasites were selected 24 h post transfection by addition of mycophenolic acid (MPA; 25µg/mL) and xanthine (XAN; 50 µg/mL) to culture medium. Lines were cloned, and successful 5’ and 3’ integration of the *mAID_Myc_HXGPRT* cassette was confirmed using primer pairs 37/38 and 39/40, respectively. Absence of WT was confirmed using primers 41 and 42.

To complement iKD CDPK1 strain with CDPK1-expressing constructs, pUPRT_CDPK1_ HA_T2A_GFP and pUPRT_CDPK1(G2A)_HA_T2A_mCherry plasmids were linearized with AclI and individually co-transfected with pSAG1_Cas9-U6_sgUPRT. Transgenic parasites were subjected to 5’-fluo-2’-deoxyuridine (FUDR) selection (5 µM) 24 h post transfection. Resistant parasites were cloned and successful 5’ and 3’ integration of was confirmed using primer pairs 49/50 and 51/52, respectively. Disruption of the endogenous *Uprt* locus was confirmed using primer pair 47/48.

### Cell culture

Parasites were cultured in Human foreskin fibroblasts (HFFs) monolayers in Dulbecco’s Modified Eagle Medium (DMEM), GlutaMAX (Thermo Fisher) supplemented with 10% heat-inactivated foetal bovine serum (FBS; Life technologies), at 37°C and 5% CO_2_. All strains and host cell lines tested negative for the presence of mycoplasma.

### Metabolic tagging and cell lysis

Upon infection of HFF monolayers the medium was removed and replaced by fresh culture media supplemented with 25 µM YnMyr (Iris Biotech) or Myr (Tokyo Chemical Industry). The parasites were then incubated for 16 h, washed with PBS (2x) and lysed on ice using a lysis buffer (PBS, 0.1% SDS, 1% Triton X-100, EDTA-free complete protease inhibitor (Roche Diagnostics)). Lysates were kept on ice for 20 min and centrifuged at 17,000 × g for 20 min to remove insoluble material. Supernatants were collected and protein concentration was determined using a BCA protein assay kit (Pierce).

### Click reaction and pull down

Lysates were thawed on ice. Proteins (100-300 µg) were taken and diluted to 1 mg/mL using the lysis buffer. A click mixture was prepared by adding reagents in the following order and by vortexing between the addition of each reagent: a capture reagent (stock solution 10 mM in water, final concentration 0.1 mM), CuSO_4_ (stock solution 50 mM in water, final concentration 1 mM), TCEP (stock solution 50 mM in water, final concentration 1 mM), TBTA (stock solution 10 mM in DMSO, final concentration 0.1 mM). Following the addition of the click mixture the samples were vortexed (room temperature, 1 h), and the reaction was stopped by addition of EDTA (final concentration 10 mM). Subsequently, proteins were precipitated (chloroform/methanol, 0.25:1, relative to the sample volume), the precipitates isolated by centrifugation (17,000 × g, 10 min), washed with methanol (1 × 400 µL) and air dried (10 min). The pellets were then resuspended (final concentration 1 mg/mL, PBS, 0.4 % SDS) and the precipitation step was repeated to remove excess of the capture reagent. Next, samples were added to 15 µL of pre-washed (0.2 % SDS in PBS (3 × 500 µL)) Dynabeads® MyOne(tm) Streptavidin C1 (Invitrogen) and gently vortexed for 90 min. The supernatant was removed and the beads were washed with 0.2 % SDS in PBS (3 × 500 µL).

### SDS-PAGE, in gel fluorescence and Western blotting (WB)

Beads were supplemented with 2% SDS in PBS (20 µL) and 4x SLB (Invitrogen), boiled (95°C, 10 min), centrifuged (1,000 × g, 2 min) and loaded on 10% or 4-20% SDS-PAGE gel (Bio-Rad). Following electrophoresis (60 min, 160V), gels were washed with water (3x). In-gel fluorescence was detected using a Pharos FX Plus Imager (Bio-Rad) and the protein loading was checked by Coomassie staining. For WB proteins were transferred (25 V, 1.3 A, 7 min) onto nitrocellulose membranes (Bio-Rad) using Bio-Rad Trans Blot Turbo Transfer system. After brief wash with PBS-T (PBS, 0.1% Tween-20) membranes were blocked (5% milk, TBS-T, 1h) and incubated with primary antibodies (5% milk, TBS-T, overnight, 4°C) at the following dilutions: rat anti-HA (1:1000; Roche Diagnostics), mouse anti-Myc (1:1000; Millipore), mouse anti-Ty1 (1:2000; Thermo Fisher), rabbit anti-Gra29 (1:1000; Moritz Treeck Lab), rabbit anti-SFP1 (1:1000; Moritz Treeck Lab), mouse anti-Toxoplasma [TP3] (1:1000; Abcam), mouse anti-CDPK1 (1:3000; John Boothroyd Lab), rabbit anti-SAG1 (1:10,000; John Boothroyd Lab), rabbit anti-GAP45 (1:1000; Peter Bradley Lab), mouse anti-GFP (1:1000, Roche Diagnostics) and rabbit anti-mCherry (1:1000, Abcam). Following washing (TBS-T, 3x) membranes were incubated with IR dye-conjugated secondary antibodies (1:10,000, 5% milk, TBS-T, 1 h) and after a final washing step imaged on a LiCOR Odyssey imaging system (LI-COR Biosciences). In case of biotin WB, membranes were blocked with 3% BSA and incubated with Streptavidin-HRP (1:4000; Thermo Scientific) in 0.3% BSA, PBS-T for 1 h. ECL Western Blotting Detection Reagent (GE Healthcare) was then used for chemiluminescence imaging on a ChemiDoc MP Imaging System (Bio-Rad).

### Synthesis of capture reagents

TEV reagent: Solid phase synthesis took place on a CF peptide synthesizer (Intavis) using a Rink Amide LL resin (100 µmol; Merck) and *N*(α)-Fmoc amino acids, including Fmoc-Lys(N_3_)-OH (Fluorochem) and Fmoc-Gly-(Dmb)Gly-OH (Merck). HCTU was used as the coupling reagent with 5-fold excess of amino acids. Fmoc-Lys(Biotin)-OH (4 eq; Merck) in 6 ml DMSO:NMP (1:1) was coupled manually after automated assembly of the rest of the chain. DIPEA (4 eq) was added, followed by HOBt (1 M, 4 eq) in NMP. After 3 min DIC (4 eq) was added, then after 30 min the solution was added to the resin and allowed to react overnight. The resin was washed with DCM and DMF prior to manual Fmoc removal and acetylation. The peptide was cleaved from the resin and protecting groups removed by addition of a cleavage solution (95% TFA, 2.5% H_2_O, 2.5% TIS). After 2 h, the resin was removed by filtration and peptides were precipitated with diethyl ether on ice. The peptide was isolated by centrifugation, then dissolved in H_2_O and freeze dried overnight. After dissolving in methanol, portions of the peptide were purified on a C8 reverse phase HPLC column (Agilent PrepHT Zorbax 300SB-C8, 21.2×250 mm, 7 m) using a linear solvent gradient of 13-50% MeCN (0.08% TFA) in H_2_O (0.08% TFA) over 40 min at a flow rate of 8 mL/min. The peak fraction was analyzed by LC–MS on an Agilent 1100 LC-MSD. The calculated molecular weight of the peptide was in agreement with the mass found. Calculated MW: 1804.08, actual mass: 1803.87.

Trypsin reagent: Solid phase synthesis took place on a CF peptide synthesizer (Intavis) using a Fmoc-PEG-Biotin NovaTag ™ resin (100 µmol; Merck), 2-Azidoacetic acid (Fluorochem) and *N*(α)-Fmoc amino acids, including Fmoc-Lys(MMT)-OH (Merck). HATU was used as the coupling reagent with 5-fold excess of amino acids. Following chain assembly, the MMT protecting group was removed from the peptidyl-resin by treatment with 1% TFA in DCM (10 mL for 2 min × 8) and the resin washed with DCM and DMF. Next 5-TAMRA (4eq; Anaspec) was dissolved in 1 ml DMSO:NMP (1:1). DIPEA (4 eq) was added, followed by HOBt (1 M, 4 eq) in NMP. After 3 min DIC (4 eq) was added, then after 30 min the solution was added to the resin and allowed to react overnight. After washing the resin with DMF and DCM, the peptide was cleaved from the resin and protecting groups removed by addition of a cleavage solution (95% TFA, 2.5% H_2_O, 2.5% TIS). After 2 h, the resin was removed by filtration and peptides were precipitated with diethyl ether on ice. The peptide was isolated by centrifugation, then dissolved in H_2_O and freeze dried overnight. After dissolving in MeCN:H_2_O (1:1), portions of the peptide were purified on a C8 reverse phase HPLC column (Agilent PrepHT Zorbax 300SB-C8, 21.2×250 mm, 7 m) using a linear solvent gradient of 10-50% MeCN (0.08% TFA) in H_2_O (0.08% TFA) over 40 min at a flow rate of 8 mL/min. The peak fraction was analyzed by LC–MS on an Agilent 1100 LC-MSD. The calculated molecular weight of the peptide was in agreement with the mass found. Calculated MW: 1396.31, actual mass: 1395.60.

### Sample preparation for MS-based proteomics

Click reaction - Reagent **1** and **2**: lysates were thawed on ice and the click reaction was carried out with 1 mg of proteins at 2 mg/mL. Proteins were captured by adding a mixture of respective capture reagent (final concentration 0.1 mM), CuSO_4_ (final concentration 1 mM), TCEP (final concentration 1 mM) and TBTA (final concentration 0.1 mM). The samples were vortex-mixed (room temperature, 1 h) before the addition of EDTA (final concentration 10 mM), methanol (4 volumes), chloroform (1 volume), and water (3 volumes). The samples were vortex-mixed briefly, centrifuged (10,000 × g, 20 min) and the resulting pellets were either washed with methanol (4 volumes) and dried (reagent **1**) or resuspended (at 2 mg/mL, 1% SDS in PBS) after which the precipitation step was repeated and the resulting pellets washed with methanol (4 volumes) and dried (reagent **2**). Reagent **3**: lysates were thawed on ice and the click reaction was carried out with 1 mg of proteins at 2 mg/mL. Proteins were captured by sequential addition of the capture reagent (final concentration 0.1 mM), TCEP (final concentration 1 mM), TBTA (stock in DMSO:t-Butanol 1:4, final concentration 0.1 mM) and CuSO_4_ (final concentration 1 mM) with mixing between each step. The samples were incubated at room temperature for 1 h before the addition of EDTA (final concentration 10 mM), methanol (4 volumes), chloroform (1 volume), and water (3 volumes). The samples were vortex-mixed briefly, centrifuged (10,000 × g, 20 min) and the resulting pellets were washed with methanol (4 volumes) and dried. Subsequently, the dried pellets were resuspended in 2% SDS in PBS and, once completely dissolved, PBS was added (final concentration 0.8% SDS, 2 mg/mL). For samples treated with base, NaOH was added (final concentration 0.2 M, 1 followed by neutralization with equivalent amount of HCl. Base treated and untreated samples were then diluted (1 mg/mL, 0.4% SDS, 100 mM DTT) before pull down.

Pull down, reduction and alkylation - NeutrAvidin agarose resin (Thermo Scientific) was washed with 0.2% SDS in PBS (3x). Typically, 50 µL of bead slurry was used for 1 mg of lysate. The samples were added to beads and the enrichment was carried out with gentle mixing (2 h, room temperature). Following the removal of supernatants, the beads were sequentially washed with 1% SDS in PBS (3x), 4 M urea in PBS (2x) and 50 mM ammonium bicarbonate (3x). The samples were reduced (5 mM DTT, 55°C, 30 min) and cysteines alkylated (10 mM iodoacetamide, room temperature, 30 min) in the dark with washing the beads (2x, 50 mM ammonium bicarbonate) after each step.

Protein digestion - for samples processed with reagent **1** and **2** as well as for supernatants (proteomes) MS grade trypsin (Promega) was used at 1:1000 w/w protease:protein, and samples were incubated overnight at 37°C. For reagent **3** two digestion strategies were used. TEV I: beads were washed (2x) with water followed by TEV buffer (50 mM TrisHCl, 0.5 mM EDTA, 1 mM DTT, pH 8.0) and the TEV protease (50 units, Invitrogen) was added. Samples were incubated overnight at 30°C. Supernatant was then removed and beads washed with TEV buffer (1x, 50 µL). The wash fraction was combined with the supernatant and stored at 4°C. A fresh portion of TEV protease (20 units) was then added to beads which were incubated for additional 6 h at 30°C. The supernatant and wash were combined with the first TEV elution. MS grade Trypsin was subsequently added at 1:1000 w/w protease:protein, and samples were incubated overnight at 37°C. TEV II: samples were incubated overnight at 37 °C with MS grade Trypsin at 1:1000 w/w protease:protein. The supernatant was removed (fraction 1) and beads washed with water and TEV buffer (2x each). The TEV protease was then added (50 units) and beads incubated overnight at 30°C (fraction 2).

Stage tip - samples were desalted prior to LC-MS/MS using Empore C18 discs (3M). Each stage tip was packed with one C18 disc, conditioned with 100 µL of 100% methanol, followed by 200 µL of 1% TFA. The samples were loaded in 1% TFA, washed 3 times with 200 µL of 1% TFA and eluted with 50 µL of 50% acetonitrile, 5% TFA. Desalted peptides were vacuum dried in preparation for LC-MS/MS analysis.

### LC-MS/MS

Samples were resuspended in 0.1% TFA and loaded on a 50 cm Easy Spray PepMap column (75 μm inner diameter, 2 μm particle size, Thermo Fisher Scientific) equipped with an integrated electrospray emitter. Reverse phase chromatography was performed using the RSLC nano U3000 (Thermo Fisher Scientific) with a binary buffer system (solvent A: 0.1% formic acid, 5% DMSO; solvent B: 80% acetonitrile, 0.1% formic acid, 5% DMSO) at a flow rate of 250 nL/min. Samples processed with reagent **1** were run on a linear gradient of 2-35% in 90 min with a total run time of 120 min including column conditioning. Samples processed with reagents **2** and **3** were run on a linear gradient of 2-40% B or 2- 55% B (TEV II myristoylated peptide fraction) in 155 min with a total run time of 180 min including column conditioning. The nanoLC was coupled to a QExactive mass spectrometer using an EasySpray nano source (Thermo Fisher Scientific). The Q-Exactive was operated in data-dependent mode, acquiring HCD MS/MS scans (R=17,500) after an MS1 survey scan (R=70, 000) on the 10 most abundant ions using MS1 target of 1E6 ions, and MS2 target of 5E4 ions. The maximum ion injection time utilized for MS2 scans was 120 ms, the HCD normalized collision energy was set at 28 and the dynamic exclusion was set at 30 seconds. The peptide match and isotope exclusion functions were enabled.

### Data analysis

Acquired raw files were processed with MaxQuant, version 1.5.2.8 (Cox and Mann, 2008) and peptides were identified from the MS/MS spectra searched against *Toxoplasma gondii* (ToxoDB) and *Homo sapiens* (UniProt) proteomes using Andromeda (Cox et al., 2011) search engine. Cysteine carbamidomethylation was selected as a fixed modification and methionine oxidation was selected as a variable modification. The enzyme specificity was set to trypsin with a maximum of 2 missed cleavages. The precursor mass tolerance was set to 20 ppm for the first search (used for mass re-calibration) and to 4.5 ppm for the main search. The datasets were filtered on posterior error probability (PEP) to achieve a 1% false discovery rate on protein, peptide and site level. Other parameters were used as pre-set in the software. “Unique and razor peptides” mode was selected to allow identification and quantification of proteins in groups (razor peptides are uniquely assigned to protein groups and not to individual proteins). Label-free quantification (LFQ) in MaxQuant was performed using a built-in label-free quantification algorithm (Cox and Mann, 2008) enabling the ‘Match between runs’ option (time window 0.7 minutes) within replicates. Each experiment comprised of replicates treated with YnMyr and the same number of replicates treated with Myr control. The LFQ is based on intensities of proteins calculated by MaxQuant from peak intensities and based on the ion currents carried by peptides whose sequences match a specific protein or a protein group to provide an approximation of abundance.

Myristoylated peptide search in MaxQuant was performed as described above applying the following variable modifications: cysteine carbamidomethylation, +463.2907 (reagent **2**) and +491.3220 (reagent **3**) at any peptide N-terminus and cysteine residues. In addition, the minimum peptide length was reduced to 6 amino acids and the ‘Match between runs’ option was disabled. MaxQuant utilizes a scoring algorithm when matching experimental MS/MS spectra with a library of theoretical spectra generated from the in silico digestion of proteins within databases selected for the search. The algorithm is used to evaluate the quality of peptide-spectrum matches (PSMs). To each PSM, MaxQuant also attributes a delta score, which is a difference between scores associated with the match to the best peptide candidate and the second best match within the database. The higher the score and the delta score, the more reliable the identification. In order to reduce a possibility for a false peptide sequence assignment even further, we applied relatively high delta score thresholds (20 vs 6 pre-set as default) for all myristoylated peptides in our analysis.

MaxQuant output files were processed with Perseus, version 1.5.0.9 (Tyanova et al., 2016) as described in the Results section and in Tables S1-S4.

### Depletion of mAID tagged CDPK1

Parasites were treated with 500 µM IAA or an equivalent volume of vehicle (ethanol) for at least 2 h prior to WB analysis.

### Depletion of *Mic7*

Parasites were treated with 50 nM rapamycin or an equivalent volume of vehicle (DMSO) for 4 h. The media was then replaced and the parasites allowed to grow for at least 24 h prior to PCR and WB analysis.

### Plaque formation

CDPK1 lines: Parasites were harvested by syringe lysis, counted, and 200 parasites were seeded on confluent HFF monolayers grown in 24-well plates (Falcon). Wells were treated with 500 µM IAA or vehicle (ethanol) and plaques were allowed to form for 5 days. MIC7 lines: Parasites were harvested by syringe lysis, counted, and 400 parasites were seeded on confluent HFF monolayers grown in 24-well plates (Falcon). Parasites were allowed to invade overnight prior to treatment with 50 nM rapamycin or vehicle (DMSO) for 4 h. Following media replacement to standard culture media, plaques were allowed to form for 5 days. iKO MIC7 line: Parasites were harvested by syringe lysis, counted, and 100 parasites were seeded on confluent HFF monolayers grown in 24-well plates (Falcon). Parasites were allowed to invade overnight prior to treatment with vehicle (DMSO) for 4 h. Following media replacement to standard culture media, plaques were allowed to form for 7 days. Plaque formation was assessed by inspecting the methanol fixed and 0.1% crystal violet stained HFF monolayers.

### Fractionation

RH Δku80Δhxgprt YFP expressing parasites were metabolically tagged with 25 µM Myr or YnMyr for 16 h. Following a PBS wash, the parasites were syringe lysed in Endo buffer (44.7 mM K_2_SO_4_, 10 mM MgSO_4_, 106 mM sucrose, 5 mM glucose, 20 mM Tris–H_2_SO_4_, 3.5 mg/ml BSA, pH 8.2) and collected by centrifugation (512 × g, 10 min). The parasites were then lysed in 300 µL of cold hypotonic buffer (10 mM HEPES, pH 7.5) supplemented with protease inhibitors (Roche), passed through 25G needle (5x) and left on ice for 40 min. Next, lysates were pelleted by centrifugation (16,000 × g, 20 min, 4 °C) and the resulting cytosolic fraction was subjected to an additional high speed (100,000 × g, 1 h, 4 °C) centrifugation step. To avoid the loss of the high speed pellet, only half of the cytosolic fraction was removed at this point. Each fraction was then taken up in 0.4% (final) SDS HEPES, clicked to a capture reagent and pulled down as described above. Myristoylation-dependent partitioning was revealed by SDS-PAGE and Western blotting.

Myristoylation-dependent fractionation for CDPK1 complemented WT and Mut lines: parasites were seeded 24 h prior experiment. Following a PBS wash, the parasites were syringe lysed in Endo buffer (44.7 mM K_2_SO_4_, 10 mM MgSO_4_, 106 mM sucrose, 5 mM glucose, 20 mM Tris–H_2_SO_4_, 3.5 mg/ml BSA, pH 8.2) and collected by centrifugation (512 × g, 10 min). The parasites were then lysed in 300 µL of cold hypotonic buffer (10 mM HEPES, pH 7.5) supplemented with protease inhibitors (Roche), passed through 25G needle (5x) and left on ice for 40 min. Next, lysates were pelleted by centrifugation (100,000 × g, 1h, 4 °C), the cytosolic fraction removed and cytosolic proteins precipitated with methanol/chloroform. Both pellet and cytosolic fractions were dissolved in 2% (final) SDS PBS and myristoylation-dependent partitioning was revealed by SDS-PAGE and Western blotting.

### Egress Assay

Parasites were added to HFF monolayer and grown for 24 h in a 96 well µplate. The wells were then treated with 500 µM IAA or an equivalent volume of vehicle (ethanol) for 2 h and then washed with PBS (2x). The media was exchanged for 100 µl Ringers solution (155 mM NaCl, 3 mM KCl, 2 mM CaCl_2_, 1 mM MgCl_2_, 3 mM NaH_2_PO_4_, 10 mM HEPES, 10 mM glucose) and the plate was placed on a heating block to maintain the temperature at 37°C. To artificially induce egress, 50 µl of Ringer’s solution containing 24 µM ionophore (8 µM final, Thermo) was added to each well. At specified time points the cells were fixed by adding 33 µl 16% formaldehyde (3% final) for 15 min. Cells were washed in PBS (3x) and stained with rabbit anti-*Tg*CAP 1:2000 (Hunt et al., bioRxiv) followed by goat anti-rabbit Alexa Fluor 488 (1:2000) and DAPI (5 µg/ml). Automated image acquisition of 25 fields per well was performed on a Cellomics Array Scan VTI HCS reader (Thermo Scientific) using a 20x objective. Image analysis was performed using the Compartmental Analysis BioApplication on HCS Studio (Thermo Scientific). Egress levels were determined in triplicate for three independent assays. Vacuole counts were normalized to t = 0 to determine how many intact vacuoles had remained after egress. The results were statistically tested using one-way ANOVA with Tukey’s multiple comparison test in GraphPad Prism® 7. The data are presented as mean ± s.d.

### Invasion assay

Parasites were treated with 50 nM rapamycin for 4 h and after replacing the media allowed to grow for 24 h. Red/green invasion assays were then performed. Parasites were lysed in an invasion non-permissive buffer, Endo buffer (44.7 mM K_2_SO_4_, 10 mM MgSO_4_, 106 mM sucrose, 5 mM glucose, 20 mM Tris–H_2_SO_4_, 3.5 mg/ml BSA, pH 8.2). 250 µl of 8×10^5^ parasites/ml in Endo buffer were added to each well of a 24-well flat-bottom plate (Falcon) containing a coverslip with a confluent HFF monolayer. The plates were spun at 129 × g for 1 min at 37 °C to deposit parasites onto the monolayer. The Endo buffer was gently removed and replaced with invasion permissive medium (1% FBS/DMEM). Parasites were allowed to invade for 15 min at 37 °C, after which the monolayer was gently washed with PBS and fixed with 3% formaldehyde for 15 min at room temperature. Extracellular (attached) parasites were stained with mouse anti-*Toxoplasma* [TP3] (Abcam) 1:1000 and goat anti-mouse Alexa Fluor 488 before permeabilization (0.2% Triton X-100/PBS) and detection of intracellular (invaded) parasites with rabbit anti-*Tg*CAP 1:2000 (Hunt et al., bioRxiv) and goat anti-rabbit Alexa Fluor 594. For each replicate, at least 5 random fields were imaged with a 40x objective. Three independent experiments were performed in duplicate. The number of intracellular (594+/488-) and extracellular (594+/488+) parasites was determined by counting, in a blinded fashion, at least 275 parasites per strain. The parasite counts in the MIC7 iKO and cMut lines were normalized to the cWT, and results were statistically tested with a one-way ANOVA with Dunnett’s multiple comparison test in GraphPad Prism® 7. The data are presented as mean ± s.d. For estimation of the parasite attachment efficiency, the number of all (594+) parasites was used and the results were statistically tested as above.

### Immunofluorescence analysis

Parasite-infected HFF monolayers grown on glass coverslips were fixed with 3% formaldehyde for 15 min prior to washing with PBS. Fixed cells were then permeabilised (PBS, 0.1% Triton X-100, 10 min), blocked (3% BSA in PBS, 1 h) and labeled with primary antibodies at the following dilutions: rat anti-HA (1:1000; Roche), mouse anti-Myc (1:1000; Millipore), mouse anti-Ty1 (1:500; Thermo Fisher), rabbit anti-MIC2 (1:5000; Vernon Carruthers Lab). Labeled proteins were visualized with Alexa Fluor-conjugated secondary goat antibodies (1:2000, Life Technologies). Nuclei were visualized with the DNA stain (DAPI; Sigma) added at 5 µg/ml with the secondary antibody. Stained coverslips were mounted on glass slides with Slowfade (Life Technologies) and imaged on a Nikon Eclipse Ti-U inverted fluorescent microscope using 100x oil objective. Images were analysed using Nikon NIS-Elements imaging software.

### MIC7 expression in tachyzoites and bradyzoites

HFF monolayers were infected with Pru Δhxgprt parasites in triplicate. For tachyzoite samples an MOI of 1 was used for a 27 h infection. For bradyzoite samples monolayers were infected at an MOI of 0.8 for 3.5 h, washed and grown in switch conditions (RPMI, 1% FBS, pH 8.1, ambient CO_2_) for 3 days. Triplicate samples were lysed in 2 mL ice cold lysis buffer (50 mM Tris-HCl, 75 mM NaCl, 8 M Urea pH 8.2), supplemented with protease (Roche Diagnostics) and phosphatase (Phos Stop, Roche Diagnostics) inhibitors. Lysis was followed by sonication to reduce sample viscosity (30% duty cycle, 3 × 30 seconds bursts, on ice). Protein concentration was measured using a BCA protein assay kit (Thermo Fisher Scientific). Lysates (1 mg per condition) were subsequently processed for mass spectrometry as described (Yang et al., 2019) and data analysis performed as explained in Table S4.

## Supplemental figure titles and legends

**Figure S1, related to Figure 1.**
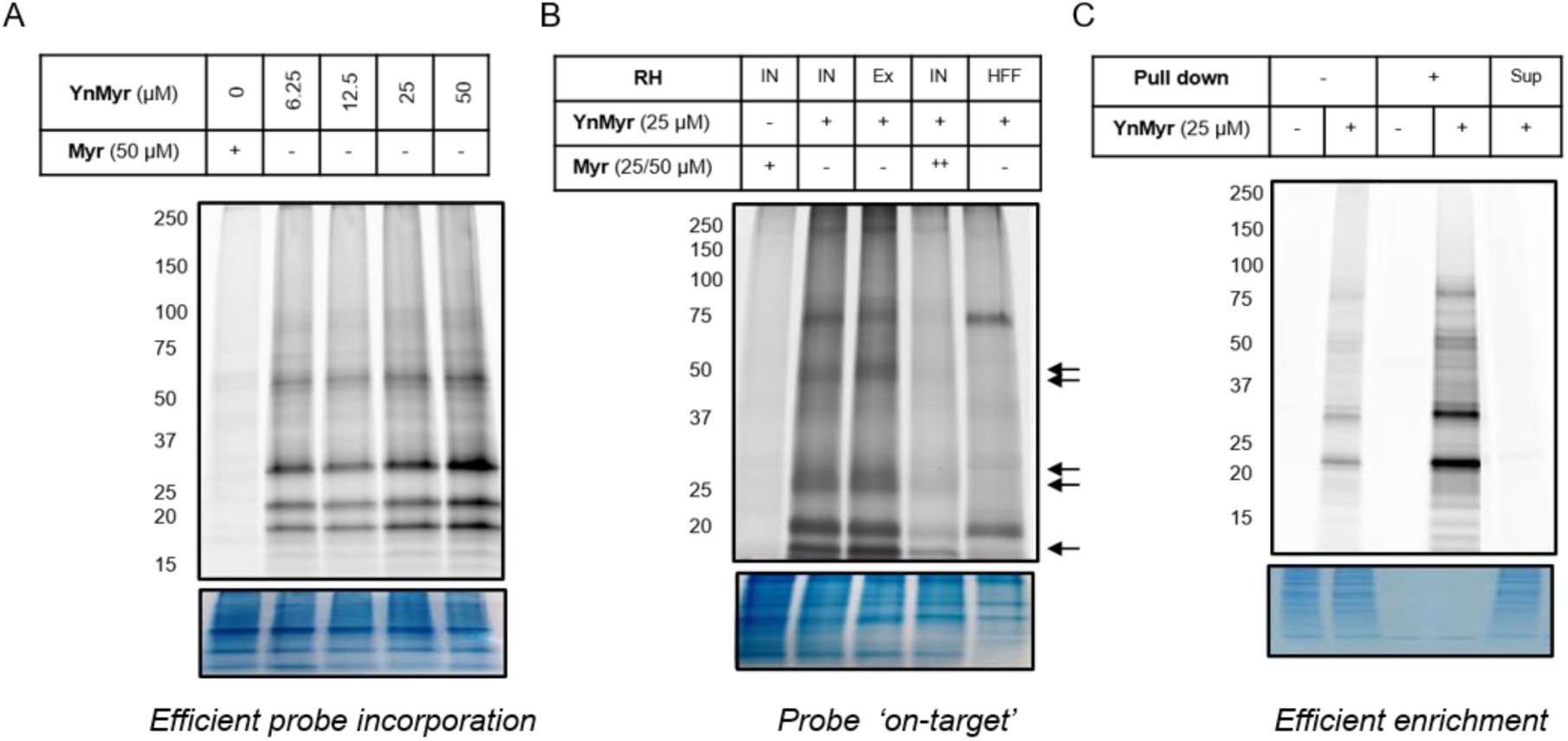
Metabolic tagging optimization. (A) In gel fluorescence imaging of protein tagging with increasing concentrations of YnMyr over a 16 h period in RH parasites. (B) In gel fluorescence visualization of protein tagging with YnMyr in intracellular (IN) and extracellular (Ex) RH parasites as well as in uninfected human foreskin fibroblasts (HFFs). Parasite-specific bands are indicated by arrows. Tagging with YnMyr is outcompeted by excess myristate (50 µM = ++). (C) In gel fluorescence analysis of YnMyr-dependent pull down efficiency, Sup = supernatant after enrichment. All bottom panels show loading control by Coomassie staining.

**Figure S2, related to Figure 2.**
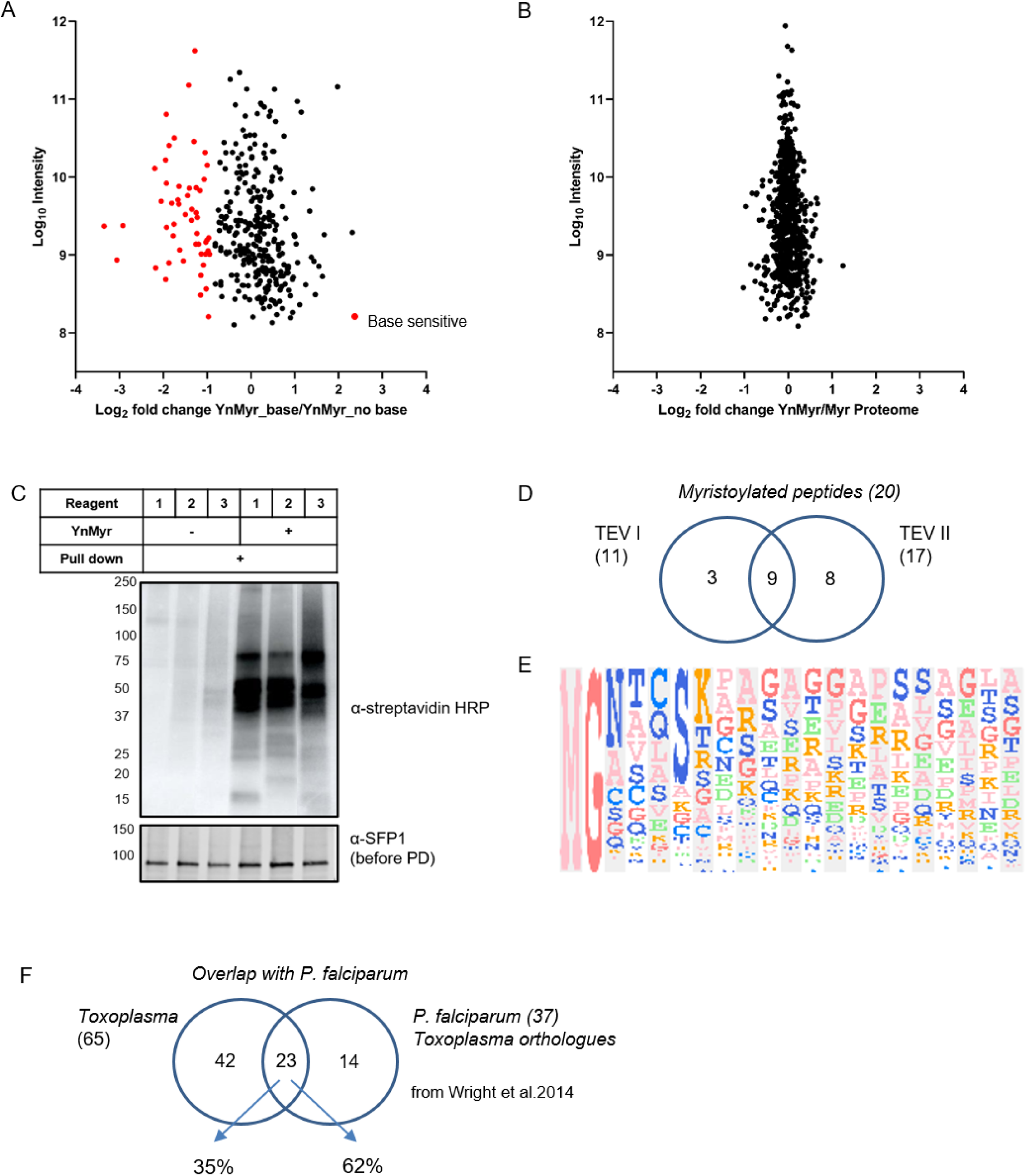
Identification of the myristoylated proteome in *T. gondii*. (A) Label free quantification of YnMyr enrichment in base-treated vs untreated samples. Proteins with log_2_ fold change < −1 are assigned as base sensitive (YnMyr incorporation through ester bonds) and are highlighted in red. See also Table S1. (B) Label free quantification of change in total protein abundance between YnMyr and Myr samples. See also Table S1. (C) Evaluation of YnMyr-dependent enrichment efficiency for capture reagents used in this study. Visualization performed by Western blotting with α- streptavidin HRP, SFP1 (TGGT1_289540) was used as loading control. (D) Venn diagram illustrating the overlap between myristoylated peptides identified with reagent **3** used in TEV I vs TEV II strategy. The number of peptides per strategy and in total is given in parenthesis. (E) Sequence logo illustrating the amino acid distribution within the 20 *N*-terminal residues of all targets. Amino acids at each position (1-20) are ordered by the frequency of occurrence. Sequence logo was created using the build-in tool within the Perseus software. (F) Venn diagram illustrating the overlap of the myristoylated proteome identified in this study with that of *P. falciparum*.

**Figure S3, related to Figure 3.**
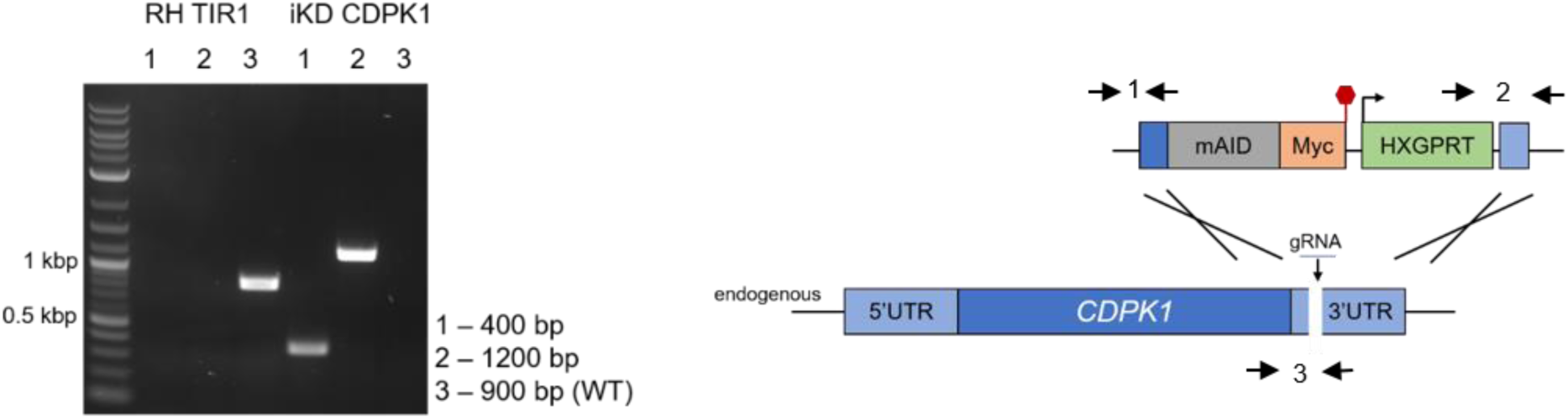
Inducible knock-down of CDPK1. PCR analysis confirming correct integration of the mAID cassette at the *C*-terminus of endogenous *cdpk1* in the iKD line. Primers are indicated by arrows. Red hexagon represents STOP codon. bp – base pairs.

**Figure S4, related to Figure 4.**
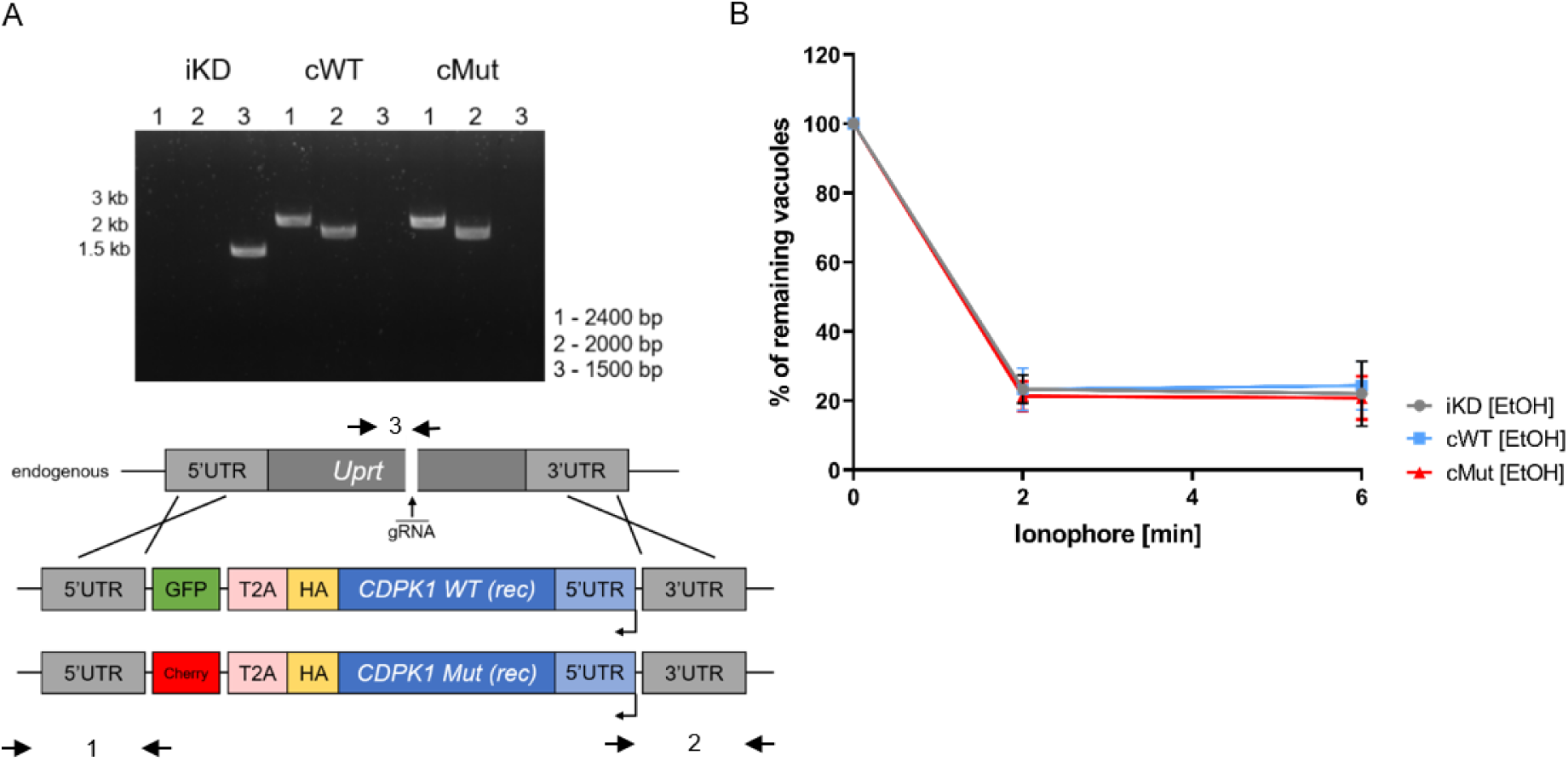
Complementation of the CDPK1 iKD line. (A) PCR analysis confirming correct integration of the complementation constructs encoding the WT and myristoylation mutant (Mut) copies of *cdpk1* at the *uprt* locus of the iKD line. Primers are indicated by arrows. bp – base pairs. (B) Both complemented lines egress from host cells within 2 min post ionophore treatment in the absence of IAA. Intracellular parasites were treated with EtOH for 2 h and egress was initiated by addition of 8 µM A23187. The number of intact vacuoles was monitored over the course of 6 min. Each data point is an average of three biological replicates, each in technical triplicate, error bars represent standard deviation.

**Figure S5, related to Figure 5.**
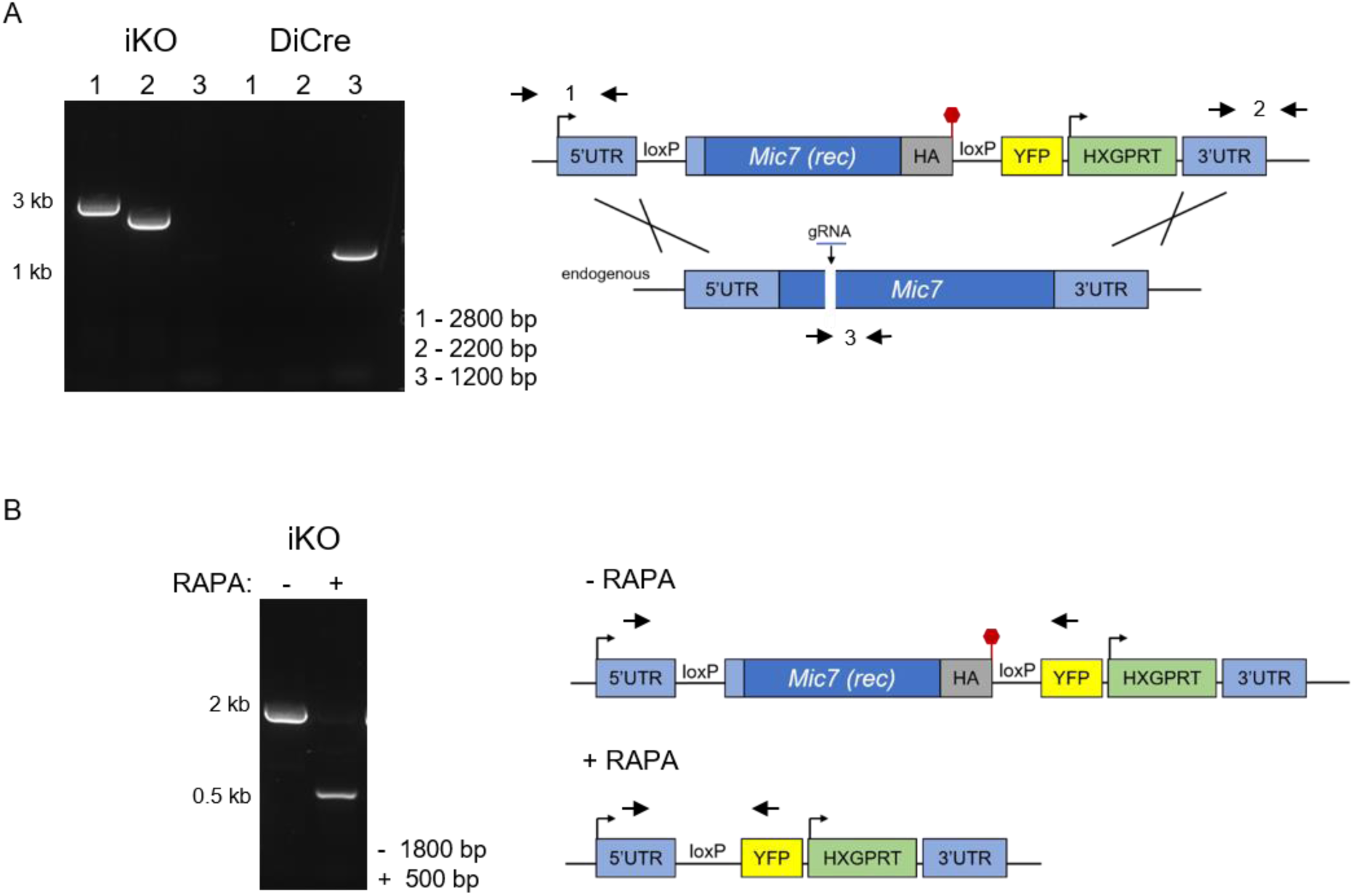
Inducible knock-out of MIC7. (A) PCR analysis confirming correct integration of the floxed and recodonized version of *mic7* in the iKO line. Primers are indicated by arrows. Red hexagon represents STOP codon. bp – base pairs. (B) PCR analysis demonstrating that addition of rapamycin (RAPA) leads to correct excision of the floxed *mic7*. Primers are indicated by arrows. Red hexagon represents STOP codon. bp – base pairs.

**Figure S6, related to Figure 6.**
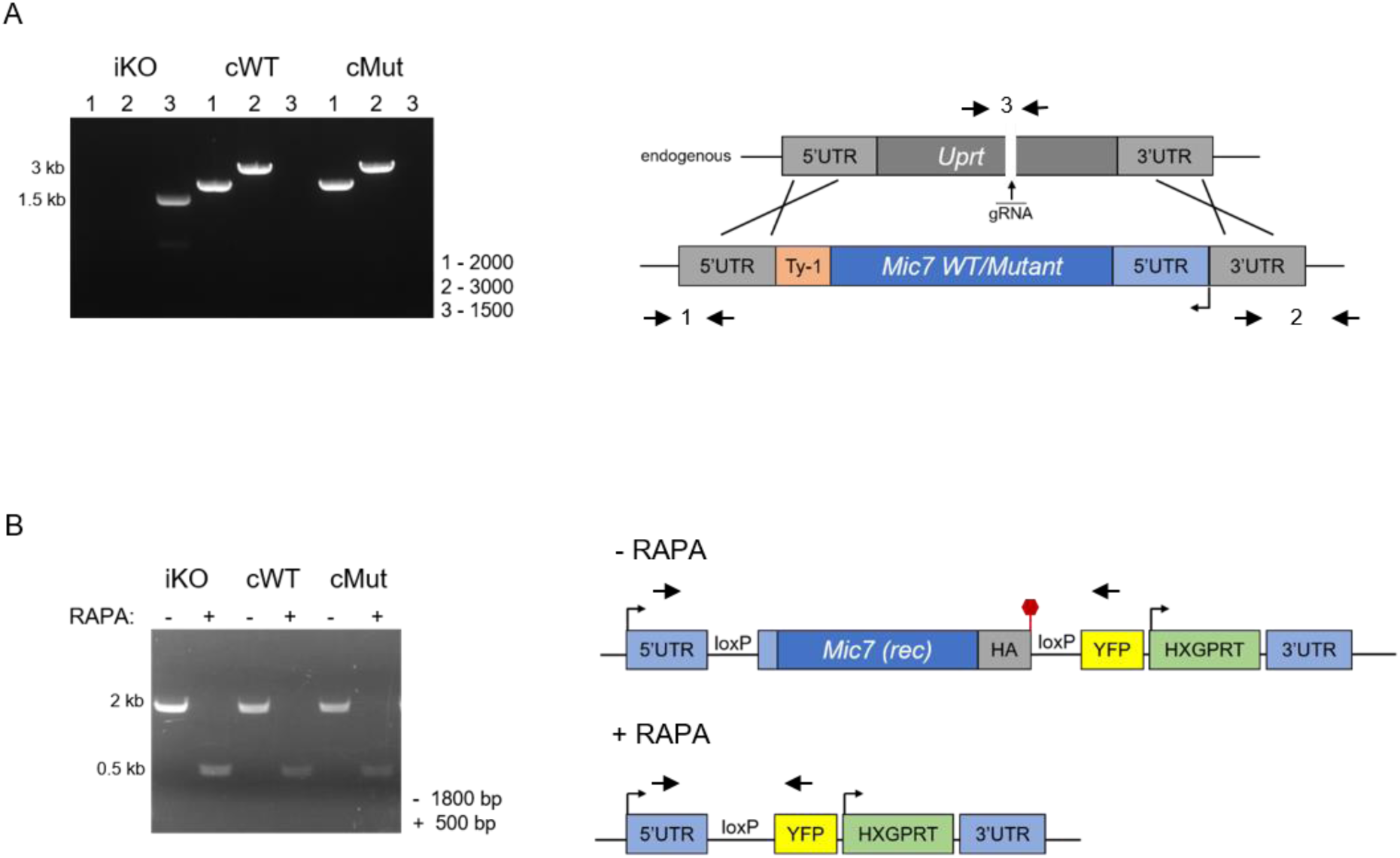
Complementation of the MIC7 iKO line. A) PCR analysis confirming correct integration of the complementation constructs encoding the WT and myristoylation mutant (Mut) copies of *mic7* at the *uprt* locus of the iKO line. Primers are indicated by arrows. bp – base pairs. (B) PCR analysis demonstrating that correct excision of the floxed *mic7* upon addition of rapamycin (RAPA) is retained in the cWT and cMut lines. Primers are indicated by arrows. Red hexagon represents STOP codon. bp – base pairs.

## Supplemental table titles and legends

**Table S1, related to Figure 2 and Figure S2. Identification of base-dependent YnMyr enrichment in *T. gondii.***

Sheet 1: *Toxoplasma* proteins with YnMyr intensities quantified irrespective of base treatment

Sheet 2: Proteins with base-sensitive enrichment

Sheet 3: MG proteins insensitive to base treatment and robustly enriched in a YnMyr-dependent manner with N_3_-biotin reagent (**1**)

Sheet 4: Analysis of proteomes (supernatants post enrichment)

**Table S2, related to Figure 2 and Figure S2. Identification of myristoylated proteins and myristoylated peptides in *T. gondii.***

Sheet 1: *Toxoplasma* proteins bearing the MG myristoylation motif

Sheet 2: Targets significantly enriched with Trypsin reagent (**2**)

Sheet 3: Targets selected based on fold change in YnMyr/Myr enrichment with TEV reagent (**3**)

Sheet 4: Myristoylated peptides found with Trypsin reagent (**2**)

Sheet 5: Myristoylated peptides found with TEV reagent (**3**)

**Table S3, related to Figure 2 and Figure S2. Myristoylated proteome of *T. gondii.***

Sheet 1: Target annotation

Sheet 2: Myristoylated proteins in *P. falciparum* and their orthologs in *Toxoplasma*

Sheet 3: Target orthologs in *P. falciparum*

**Table S4, related to Figure 5. MIC7 expression in tachyzoites and bradyzoites.**

**Table S5. Primers used for plasmid and parasite strain generation.**

## References

Alonso, A.M., Coceres, V.M., De Napoli, M.G., Nieto Guil, A.F., Angel, S.O., and Corvi, M.M. (2012). Protein palmitoylation inhibition by 2-bromopalmitate alters gliding, host cell invasion and parasite morphology in Toxoplasma gondii. Mol Biochem Parasitol 184, 39–43.

Alonso, A.M., Turowski, V.R., Ruiz, D.M., Orelo, B.D., Moresco, J.J., Yates, J.R., 3rd, and Corvi, M.M. (2019). Exploring protein myristoylation in Toxoplasma gondii. Exp Parasitol 203, 8–18.

Andenmatten, N., Egarter, S., Jackson, A.J., Jullien, N., Herman, J.P., and Meissner, M. (2013). Conditional genome engineering in Toxoplasma gondii uncovers alternative invasion mechanisms. Nat Methods 10, 125–127.

Beck, J.R., Rodriguez-Fernandez, I.A., de Leon, J.C., Huynh, M.H., Carruthers, V.B., Morrissette, N.S., and Bradley, P.J. (2010). A novel family of Toxoplasma IMC proteins displays a hierarchical organization and functions in coordinating parasite division. PLoS Pathog 6, e1001094.

Black, M.W., and Boothroyd, J.C. (2000). Lytic cycle of Toxoplasma gondii. Microbiol Mol Biol Rev 64, 607–623.

Boutin, J.A. (1997). Myristoylation. Cell Signal 9, 15–35.

Broncel, M., Serwa, R.A., Ciepla, P., Krause, E., Dallman, M.J., Magee, A.I., and Tate, E.W. (2015). Multifunctional reagents for quantitative proteome-wide analysis of protein modification in human cells and dynamic profiling of protein lipidation during vertebrate development. Angew Chem Int Ed Engl 54, 5948–5951.

Brown, K.M., Long, S., and Sibley, L.D. (2017). Plasma Membrane Association by N-Acylation Governs PKG Function in Toxoplasma gondii. MBio 8.

Brown, K.M., Long, S., and Sibley, L.D. (2018). Conditional Knockdown of Proteins Using Auxin-inducible Degron (AID) Fusions in Toxoplasma gondii. Bio Protoc 8.

Caballero, M.C., Alonso, A.M., Deng, B., Attias, M., de Souza, W., and Corvi, M.M. (2016). Identification of new palmitoylated proteins in Toxoplasma gondii. Biochim Biophys Acta 1864, 400–408.

Caffaro, C.E., Koshy, A.A., Liu, L., Zeiner, G.M., Hirschberg, C.B., and Boothroyd, J.C. (2013). A nucleotide sugar transporter involved in glycosylation of the Toxoplasma tissue cyst wall is required for efficient persistence of bradyzoites. PLoS Pathog 9, e1003331.

Chow, M., Newman, J.F., Filman, D., Hogle, J.M., Rowlands, D.J., and Brown, F. (1987). Myristylation of picornavirus capsid protein VP4 and its structural significance. Nature 327, 482–486.

Cox, J., and Mann, M. (2008). MaxQuant enables high peptide identification rates, individualized p.p.b.-range mass accuracies and proteome-wide protein quantification. Nat Biotechnol 26, 1367–1372.

Cox, J., Neuhauser, N., Michalski, A., Scheltema, R.A., Olsen, J.V., and Mann, M. (2011). Andromeda: a peptide search engine integrated into the MaxQuant environment. J Proteome Res 10, 1794–1805.

Devadas, B., Zupec, M.E., Freeman, S.K., Brown, D.L., Nagarajan, S., Sikorski, J.A., McWherter, C.A., Getman, D.P., and Gordon, J.I. (1995). Design and syntheses of potent and selective dipeptide inhibitors of Candida albicans myristoyl-CoA:protein N-myristoyltransferase. J Med Chem 38, 1837–1840.

Foe, I.T., Child, M.A., Majmudar, J.D., Krishnamurthy, S., van der Linden, W.A., Ward, G.E., Martin, B.R., and Bogyo, M. (2015). Global Analysis of Palmitoylated Proteins in Toxoplasma gondii. Cell Host Microbe 18, 501–511.

Frearson, J.A., Brand, S., McElroy, S.P., Cleghorn, L.A., Smid, O., Stojanovski, L., Price, H.P., Guther, M.L., Torrie, L.S., Robinson, D.A., et al. (2010). N-myristoyltransferase inhibitors as new leads to treat sleeping sickness. Nature 464, 728–732.

Frenal, K., Kemp, L.E., and Soldati-Favre, D. (2014). Emerging roles for protein S-palmitoylation in Toxoplasma biology. Int J Parasitol 44, 121–131.

Frenal, K., Polonais, V., Marq, J.B., Stratmann, R., Limenitakis, J., and Soldati-Favre, D. (2010). Functional dissection of the apicomplexan glideosome molecular architecture. Cell Host Microbe 8, 343–357.

Gaji, R.Y., Johnson, D.E., Treeck, M., Wang, M., Hudmon, A., and Arrizabalaga, G. (2015). Phosphorylation of a Myosin Motor by TgCDPK3 Facilitates Rapid Initiation of Motility during Toxoplasma gondii egress. PLoS Pathog 11, e1005268.

Garrison, E., Treeck, M., Ehret, E., Butz, H., Garbuz, T., Oswald, B.P., Settles, M., Boothroyd, J., and Arrizabalaga, G. (2012). A forward genetic screen reveals that calcium-dependent protein kinase 3 regulates egress in Toxoplasma. PLoS Pathog 8, e1003049.

Goldberg, J. (1998). Structural basis for activation of ARF GTPase: mechanisms of guanine nucleotide exchange and GTP-myristoyl switching. Cell 95, 237–248.

Gordon, J.I., Duronio, R.J., Rudnick, D.A., Adams, S.P., and Gokel, G.W. (1991). Protein N-myristoylation. J Biol Chem 266, 8647–8650.

Heal, W.P., Wright, M.H., Thinon, E., and Tate, E.W. (2011). Multifunctional protein labeling via enzymatic N-terminal tagging and elaboration by click chemistry. Nat Protoc 7, 105–117.

Hunt, A., Wagener, J., Kent, R., Carmeille, R., Russell, M., Peddie, C., Collinson, L., Heaslip, A., Ward, G.E., Treeck, M. (2019). Differential requirements of cyclase associated protein (CAP) for actin turnover during the lytic cycle of Toxoplasma gondii. bioRxiv; doi: https://doiorg/101101/569368.

Hutton, J.A., Goncalves, V., Brannigan, J.A., Paape, D., Wright, M.H., Waugh, T.M., Roberts, S.M., Bell, A.S., Wilkinson, A.J., Smith, D.F., et al. (2014). Structure-based design of potent and selective Leishmania N-myristoyltransferase inhibitors. J Med Chem 57, 8664–8670.

Huynh, M.H., Rabenau, K.E., Harper, J.M., Beatty, W.L., Sibley, L.D., and Carruthers, V.B. (2003). Rapid invasion of host cells by Toxoplasma requires secretion of the MIC2-M2AP adhesive protein complex. EMBO J 22, 2082–2090.

Jacot, D., and Soldati-Favre, D. (2012). Does protein phosphorylation govern host cell entry and egress by the Apicomplexa? Int J Med Microbiol 302, 195–202.

Jia, Y., Marq, J.B., Bisio, H., Jacot, D., Mueller, C., Yu, L., Choudhary, J., Brochet, M., and Soldati-Favre, D. (2017). Crosstalk between PKA and PKG controls pH-dependent host cell egress of Toxoplasma gondii. EMBO J 36, 3250–3267.

Kremer, K., Kamin, D., Rittweger, E., Wilkes, J., Flammer, H., Mahler, S., Heng, J., Tonkin, C.J., Langsley, G., Hell, S.W., et al. (2013). An overexpression screen of Toxoplasma gondii Rab-GTPases reveals distinct transport routes to the micronemes. PLoS Pathog 9, e1003213.

Lamarque, M.H., Roques, M., Kong-Hap, M., Tonkin, M.L., Rugarabamu, G., Marq, J.B., Penarete-Vargas, D.M., Boulanger, M.J., Soldati-Favre, D., and Lebrun, M. (2014). Plasticity and redundancy among AMA-RON pairs ensure host cell entry of Toxoplasma parasites. Nat Commun 5, 4098.

Liendo, A., Stedman, T.T., Ngo, H.M., Chaturvedi, S., Hoppe, H.C., and Joiner, K.A. (2001). Toxoplasma gondii ADP-ribosylation factor 1 mediates enhanced release of constitutively secreted dense granule proteins. J Biol Chem 276, 18272–18281.

Lourido, S., Jeschke, G.R., Turk, B.E., and Sibley, L.D. (2013). Exploiting the unique ATP-binding pocket of toxoplasma calcium-dependent protein kinase 1 to identify its substrates. ACS Chem Biol 8, 1155–1162.

Lourido, S., Shuman, J., Zhang, C., Shokat, K.M., Hui, R., and Sibley, L.D. (2010). Calcium-dependent protein kinase 1 is an essential regulator of exocytosis in Toxoplasma. Nature 465, 359–362.

Lourido, S., Tang, K., and Sibley, L.D. (2012). Distinct signalling pathways control Toxoplasma egress and host-cell invasion. EMBO J 31, 4524–4534.

Martin, D.D., Beauchamp, E., and Berthiaume, L.G. (2011). Post-translational myristoylation: Fat matters in cellular life and death. Biochimie 93, 18–31.

Maurer-Stroh, S., and Eisenhaber, F. (2004). Myristoylation of viral and bacterial proteins. Trends Microbiol 12, 178–185.

McCoy, J.M., Whitehead, L., van Dooren, G.G., and Tonkin, C.J. (2012). TgCDPK3 regulates calcium-dependent egress of Toxoplasma gondii from host cells. PLoS Pathog 8, e1003066.

Meissner, M., Reiss, M., Viebig, N., Carruthers, V.B., Toursel, C., Tomavo, S., Ajioka, J.W., and Soldati, D. (2002). A family of transmembrane microneme proteins of Toxoplasma gondii contain EGF-like domains and function as escorters. J Cell Sci 115, 563–574.

Mueller, C., Klages, N., Jacot, D., Santos, J.M., Cabrera, A., Gilberger, T.W., Dubremetz, J.F., and Soldati-Favre, D. (2013). The Toxoplasma protein ARO mediates the apical positioning of rhoptry organelles, a prerequisite for host cell invasion. Cell Host Microbe 13, 289–301.

Nagarajan, S.R., Devadas, B., Zupec, M.E., Freeman, S.K., Brown, D.L., Lu, H.F., Mehta, P.P., Kishore, N.S., McWherter, C.A., Getman, D.P., et al. (1997). Conformationally constrained [p-(omega-aminoalkyl)phenacetyl]-L-seryl-L-lysyl dipeptide amides as potent peptidomimetic inhibitors of Candida albicans and human myristoyl-CoA:protein N-myristoyl transferase. J Med Chem 40, 1422–1438.

Ojo, K.K., Larson, E.T., Keyloun, K.R., Castaneda, L.J., Derocher, A.E., Inampudi, K.K., Kim, J.E., Arakaki, T.L., Murphy, R.C., Zhang, L., et al. (2010). Toxoplasma gondii calcium-dependent protein kinase 1 is a target for selective kinase inhibitors. Nat Struct Mol Biol 17, 602–607.

Pomel, S., Luk, F.C., and Beckers, C.J. (2008). Host cell egress and invasion induce marked relocations of glycolytic enzymes in Toxoplasma gondii tachyzoites. PLoS Pathog 4, e1000188.

Reese, M.L., Zeiner, G.M., Saeij, J.P., Boothroyd, J.C., and Boyle, J.P. (2011). Polymorphic family of injected pseudokinases is paramount in Toxoplasma virulence. Proc Natl Acad Sci U S A 108, 9625–9630.

Robert-Gangneux, F., and Darde, M.L. (2012). Epidemiology of and diagnostic strategies for toxoplasmosis. Clin Microbiol Rev 25, 264–296.

Schlott, A.C., Mayclin, S., Reers, A.R., Coburn-Flynn, O., Bell, A.S., Green, J., Knuepfer, E., Charter, D., Bonnert, R., Campo, B., et al. (2019). Structure-Guided Identification of Resistance Breaking Antimalarial NMyristoyltransferase Inhibitors. Cell Chem Biol.

Shen, B., Brown, K.M., Lee, T.D., and Sibley, L.D. (2014). Efficient gene disruption in diverse strains of Toxoplasma gondii using CRISPR/CAS9. MBio 5, e01114–01114.

Silmon de Monerri, N.C., Yakubu, R.R., Chen, A.L., Bradley, P.J., Nieves, E., Weiss, L.M., and Kim, K. (2015). The Ubiquitin Proteome of Toxoplasma gondii Reveals Roles for Protein Ubiquitination in Cell-Cycle Transitions. Cell Host Microbe 18, 621–633.

Simons, J., Rogove, A., Moscufo, N., Reynolds, C., and Chow, M. (1993). Efficient analysis of nonviable poliovirus capsid mutants. J Virol 67, 1734–1738.

Soldati, D., and Boothroyd, J.C. (1993). Transient transfection and expression in the obligate intracellular parasite Toxoplasma gondii. Science 260, 349–352.

Soldati, D., Dubremetz, J.F., and Lebrun, M. (2001). Microneme proteins: structural and functional requirements to promote adhesion and invasion by the apicomplexan parasite Toxoplasma gondii. Int J Parasitol 31, 1293–1302.

Speers, A.E., and Cravatt, B.F. (2005). A tandem orthogonal proteolysis strategy for high-content chemical proteomics. J Am Chem Soc 127, 10018–10019.

Thinon, E., Serwa, R.A., Broncel, M., Brannigan, J.A., Brassat, U., Wright, M.H., Heal, W.P., Wilkinson, A.J., Mann, D.J., and Tate, E.W. (2014). Global profiling of co-and post-translationally N-myristoylated proteomes in human cells. Nat Commun 5, 4919.

Tosetti, N., Dos Santos Pacheco, N., Soldati-Favre, D., and Jacot, D. (2019). Three F-actin assembly centers regulate organelle inheritance, cell-cell communication and motility in Toxoplasma gondii. Elife 8.

Treeck, M., Sanders, J.L., Gaji, R.Y., LaFavers, K.A., Child, M.A., Arrizabalaga, G., Elias, J.E., and Boothroyd, J.C. (2014). The calcium-dependent protein kinase 3 of toxoplasma influences basal calcium levels and functions beyond egress as revealed by quantitative phosphoproteome analysis. PLoS Pathog 10, e1004197.

Tu, V., Mayoral, J., Sugi, T., Tomita, T., Han, B., Ma, Y.F., and Weiss, L.M. (2019). Enrichment and Proteomic Characterization of the Cyst Wall from In Vitro Toxoplasma gondii Cysts. MBio 10.

Tyanova, S., Temu, T., Sinitcyn, P., Carlson, A., Hein, M.Y., Geiger, T., Mann, M., and Cox, J. (2016). The Perseus computational platform for comprehensive analysis of (prote)omics data. Nat Methods 13, 731–740.

Uboldi, A.D., Wilde, M.L., McRae, E.A., Stewart, R.J., Dagley, L.F., Yang, L., Katris, N.J., Hapuarachchi, S.V., Coffey, M.J., Lehane, A.M., et al. (2018). Protein kinase A negatively regulates Ca2+ signalling in Toxoplasma gondii. PLoS Biol 16, e2005642.

Wallbank, B.A., Dominicus, C.S., Broncel, M., Legrave, N., Kelly, G., MacRae, J.I., Staines, H.M., and Treeck, M. (2019). Characterisation of the Toxoplasma gondii tyrosine transporter and its phosphorylation by the calcium-dependent protein kinase 3. Mol Microbiol 111, 1167–1181.

Wright, M.H., Clough, B., Rackham, M.D., Rangachari, K., Brannigan, J.A., Grainger, M., Moss, D.K., Bottrill, A.R., Heal, W.P., Broncel, M., et al. (2014). Validation of N-myristoyltransferase as an antimalarial drug target using an integrated chemical biology approach. Nat Chem 6, 112–121.

Wright, M.H., Heal, W.P., Mann, D.J., and Tate, E.W. (2010). Protein myristoylation in health and disease. J Chem Biol 3, 19–35.

Wright, M.H., Paape, D., Price, H.P., Smith, D.F., and Tate, E.W. (2016). Global Profiling and Inhibition of Protein Lipidation in Vector and Host Stages of the Sleeping Sickness Parasite Trypanosoma brucei. ACS Infect Dis 2, 427–441.

Wright, M.H., Paape, D., Storck, E.M., Serwa, R.A., Smith, D.F., and Tate, E.W. (2015). Global analysis of protein N-myristoylation and exploration of N-myristoyltransferase as a drug target in the neglected human pathogen Leishmania donovani. Chem Biol 22, 342–354.

Yang, C., Broncel, M., Dominicus, C., Sampson, E., Blakely, W.J., Treeck, M., and Arrizabalaga, G. (2019). A plasma membrane localized protein phosphatase in Toxoplasma gondii, PPM5C, regulates attachment to host cells. Sci Rep 9, 5924.

